# WDR90 is a centriolar microtubule wall protein important for centriole architecture integrity

**DOI:** 10.1101/2020.02.15.950444

**Authors:** Emmanuelle Steib, Davide Gambarotto, Marine Laporte, Natacha Olieric, Céline Zheng, Susanne Borgers, Vincent Olieric, Maeva Le Guennec, France Koll, Anne-Marie Tassin, Michel O. Steinmetz, Paul Guichard, Virginie Hamel

**Affiliations:** University of Geneva, Department of Cell Biology, Sciences III, Geneva, Switzerland; Laboratory of Biomolecular Research, Division of Biology and Chemistry, Paul Scherrer Institut, Villigen, Switzerland; Swiss Light Source, Paul Scherrer Institut, 5232 Villigen, Switzerland; Institute for Integrative Biology of the Cell (I2BC), CEA, CNRS, Univ. Paris Sud, Université Paris-Saclay, 1 Avenue de la Terrasse, 91198 Gif sur Yvette, France; Biozentrum, University of Basel, 4056 Basel, Switzerland

**Keywords:** centriole, inner scaffold, *Chlamydomonas*, human cells, Ultrastructure Expansion Microscopy, microtubules

## Abstract

Centrioles are characterized by a nine-fold arrangement of long-lived microtubule triplets that are held together by an inner protein scaffold. These structurally robust organelles experience strenuous cellular processes such as cell division or ciliary beating while performing their function. However, the molecular mechanisms underlying the stability of microtubule triplets, as well as centriole architectural integrity remain poorly understood. Here, using ultrastructure expansion microscopy (U-ExM) for nanoscale protein mapping, we reveal that POC16 and its human homolog WDR90 are components of the centriolar microtubule wall along the central core region of the centriole. We further found that WDR90 is an evolutionary microtubule associated protein with a predicted structurally homology with the ciliary inner junction protein FAP20. Finally, we demonstrate that WDR90 depletion impairs the localization of inner scaffold components, leading to centriole structural abnormalities in both human and *Chlamydomonas* cells. Altogether, this work highlights that POC16/WDR90 is a crucial evolutionary conserved molecular player participating in centriole architecture integrity.

## Introduction

Centrioles and basal bodies (referred to as centrioles from here onwards for simplicity) are conserved organelles important for the formation of the centrosome as well as for templating cilia and flagella assembly (Bornens, 2012; Breslow and Holland, 2019; Conduit et al., 2015; Ishikawa and Marshall, 2011). Consequently, defects in centriole assembly, size, structure and number lead to abnormal mitosis or defective ciliogenesis and have been associated with several human pathologies such as ciliopathies and cancer (Gönczy, 2015; Nigg and Holland, 2018; Nigg and Raff, 2009). For instance, centriole amplification, a hallmark of cancer cells, can result from centriole fragmentation in defective, over-elongated centrioles (Marteil et al., 2018).

Centrioles are characterized by a nine-fold radial arrangement of microtubule triplets, are polarized along their long axis, and can be divided in three distinct regions termed proximal end, central core and distal tip (Hamel et al., 2017). Each region displays specific structural features such as the cartwheel on the proximal end, which is crucial for centriole assembly (Nakazawa et al., 2007; Strnad et al., 2007) or the distal appendages at the very distal region, essential for membrane docking during ciliogenesis (Tanos et al., 2013). The central core region of the centriole is defined by the presence of a circular inner scaffold thought to maintain the integrity of microtubule triplets under compressive forces (Le Guennec et al., 2020). Using cryo-tomography, we recently showed that the inner centriole scaffold forms an extended helix covering ∼70% of the centriole length and that is rooted at the inner junction between the A and B microtubules (Figure 1A, B). This connection consists of a stem attaching the neighboring A and B microtubules and three arms extending from the same stem toward the centriolar lumen (Le Guennec et al., 2020) (Figure 1A, B). The stem of the inner scaffold has been detected in *Paramecium tetraurelia*, *Chlamydomonas reinhardtii* and human centrioles, suggesting that it represents an evolutionary conserved structural feature.

**Figure 1.**
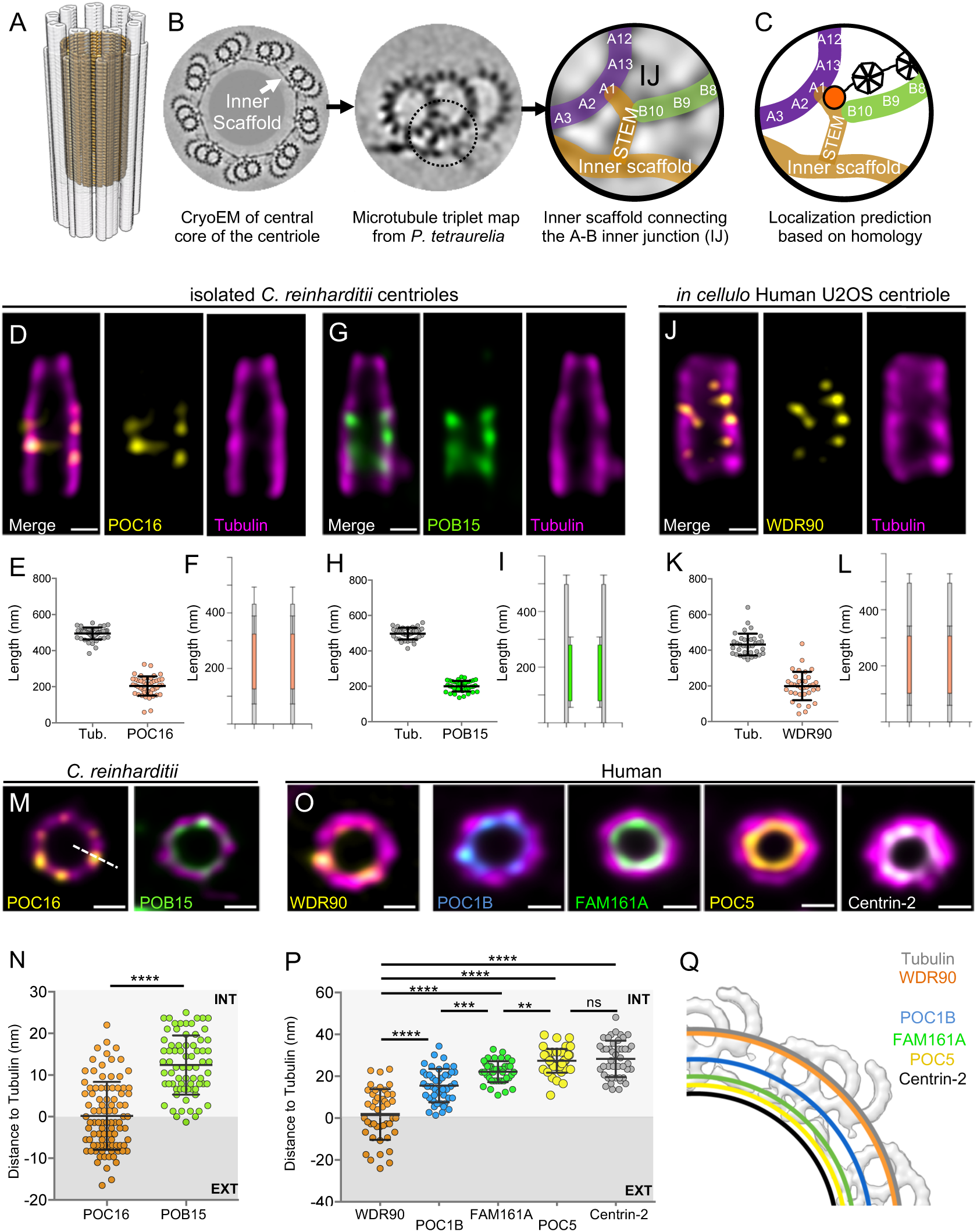
**POC16/WDR90 is a conserved central core microtubule wall component.** (A) 3D representation of a centriole highlighting the centriolar microtubule wall in light grey and the inner scaffold in yellow. (B) Cryo-EM image of the central core of *Paramecium tetraurelia* centrioles from which a microtubule triplet map has been generated (Le Guennec et al., 2020). Schematic representation of the inner junction (IJ) between A- and B-microtubules connecting the inner scaffold. (C) Schematic localization of POC16/WDR90 proteins within the IJ based on its homology to FAP20. Purple: A-microtubule, green: B microtubule, yellow: inner scaffold and stem, orange: DUF667 domain positioned at the IJ. (D) Isolated U-ExM expanded *Chlamydomonas* centriole stained for POC16 (yellow) and tubulin (magenta), lateral view. Scale bar: 100nm. (E) Respective lengths of tubulin and POC16 based on D. Average +/- SD: Tubulin: 495nm +/- 33, POC16: 204nm +/- 53, n=46 centrioles from 3 independent experiments. (F) POC16 length coverage and positioning: 41% +/- 11, n=46 centrioles from 3 independent experiments. (G) Isolated U-ExM expanded *Chlamydomonas* centriole stained for POB15 (green) and tubulin (magenta), lateral view. (H) Respective length of tubulin and POB15 based on G. Average +/- SD: tubulin= 497nm +/- 33, POB15= 200nm +/- 30, n=39 centrioles from 3 independent experiments. (I) POB15 length coverage and positioning: 40% +/- 6, n=39 centrioles from 3 independent experiments. (J) *In cellulo* U-ExM expanded human U2OS centriole stained for WDR90 (yellow) and tubulin (magenta), lateral views. (K) Respective lengths of tubulin and WDR90 based on J. Average +/- SD: Tubulin: 432nm +/- 62, WDR90: 200nm +/- 80, n=35 from 3 independent experiments. (L) WDR90 length coverage and positioning: 46% +/- 17, n=35 from 3 independent experiments. (M) Isolated U-ExM expanded *Chlamydomonas* centriole stained for tubulin (magenta) and POC16 (yellow) or POB15 (green), top view. Scale bar: 100nm. (N) Distance between the maximal intensity of tubulin and the maximal intensity of POC16 (orange) or POB15 (green) based on M. Average +/- SD: POC16= 0nm +/- 8, POB15= 12nm +/- 7. n>75 measurements/condition from 30 centrioles from 3 independent experiments. EXT: exterior or the centriole, INT: interior. (O) *In cellulo* U-ExM expanded human U2OS centriole stained for WDR90 (yellow) and tubulin (magenta), top views, or for core proteins POC1B (blue), FAM161A (green), POC5 (yellow) or Centrin-2 (white). Data set from (Le Guennec et al., 2020). Scale bare: 100nm. (P) Distance between the maximal intensity of tubulin and the maximal intensity of WDR90 (orange) or POC1B (blue), FAM161A (green), POC5 (yellow) or Centrin-2 (grey) based on O. Average +/- SD: WDR90= 2nm +/- 12, POC1B= 15nm+/- 8, FAM161A= 22nm+/-5, POC5= 27nm +/- 6 and Centrin-2= 28nm+/-9. n=45 measurements/condition from 15 to 30 centrioles from 3 independent experiments. Statistics by one-was ANOVA followed by Holm Sidak (Q) Position of WDR90 relative to the four inner scaffold components placed on the cryo-EM map of the *Paramecium* central core region (top view) (adapted from (Le Guennec et al., 2020)).

The molecular identity of some components of the inner scaffold has been uncovered using Ultrastructure Expansion Microscopy (U-ExM), which allows nanoscale localization of proteins within structural elements (Gambarotto et al., 2019). Notably, the centriolar proteins POC1B, FAM161A, POC5 and Centrin-2 have been shown to localize to the inner scaffold along the microtubule blades in human cells (Le Guennec et al., 2020). Moreover, these proteins form a complex that can bind to microtubules through the microtubule-binding protein FAM161A (Le Guennec et al., 2020; Zach et al., 2012). Importantly, a subset of these proteins has been shown to be important, such as POC5 for centriole elongation (Azimzadeh et al., 2009) as well as POC1B for centriole and basal body integrity (Pearson et al., 2009; Venoux et al., 2013). This observation highlights the role of the inner scaffold structure in providing stability to the entire centriolar microtubule wall organization. However, the exact contribution of the inner scaffold to microtubule triplets stability and how the inner scaffold is connected to the microtubule blade is unknown.

We recently identified the conserved proteins POC16/WDR90 as proteins localizing to the central core region in both *Chlamydomonas reinhardtii* and human centrioles (Hamel et al., 2017). Impairing POC16 or WDR90 functions has been found to affect ciliogenesis, suggesting that POC16/WDR90 may stabilize the microtubule wall, thereby ensuring proper flagellum or cilium assembly (Hamel et al, 2017). Interestingly, POC16 has been proposed to be at the inner junction between the A and B microtubules (H. Yanagisawa et al., 2014) through its sequence homology with FAP20, an axonemal microtubule doublet inner junction protein of *Chlamydomonas reinhardtii* flagella (Dymek et al., 2019; Ma et al., 2019; Owa et al., 2019; H. A. Yanagisawa et al., 2014). As the stem connects the A- and B-microtubules interface, these observations suggest that POC16/WDR90 may connect the inner scaffold to the microtubule triplet through this stem structure (Figure 1C), thus ensuring integrity of the centriole architecture.

In this study, using a combination of cell biology, biochemistry and Ultrastructure Expansion Microscopy (U-ExM) approaches, we establish that the conserved POC16/WDR90 proteins localize on the centriolar microtubule wall in the central core region of both *Chlamydomonas* and human cells. We further demonstrate that WDR90 is a microtubule-binding protein and that loss of this protein impairs the localization of inner scaffold components and leads to slight centriole elongation, impairment of the canonical circular shape of centrioles as well as defects in centriolar architecture integrity.

## Results

### POC16/WDR90 is a conserved microtubule wall component of the central core region

To test the hypothesis that POC16/WDR90 is a microtubule triplet component, we analyzed its distribution using U-ExM that allows nanoscale mapping of proteins inside the centriole (Gambarotto et al., 2019; Le Guennec et al., 2020). We observed first in *Chlamydomonas reinhardtii* isolated centrioles that the endogenous POC16 longitudinal fluorescence signal is restricted to the central core region as compared to the tubulin signal, which depicts total centriolar length (Figure 1D-F). From top viewed centrioles, we measured both POC16 and tubulin maximal intensity signal from the exterior to the interior of the centriole and quantified the distance between x-values (Figure 1M, N, average distance between POC16 and tubulin Δ = 0 nm +/- 8). We concluded that POC16 localizes precisely on the microtubule wall in the central core region of *Chlamydomonas* centrioles. As a control, we could recapitulate the internal localization along the microtubule wall of POB15, another central core protein (Figure 1G-I and Figure 1M, N, average distance between POB15 and tubulin Δ = 12 nm +/- 7) as previously reported using immunogold-labeling (Hamel et al., 2017). In human centrioles, the POC16 human homolog WDR90 localizes similarly to POC16 on the centriolar microtubule wall, demonstrating the evolutionary conserved restricted localization of POC16/WDR90 on microtubule triplets in the central core region of centrioles (Figure 1J-L). Of note, POC16 and WDR90 display a punctate distribution that we hypothesize to be due to the poor quality of the antibody.

Next, we compared the relative position of WDR90 to previously described inner scaffold components (Figure 1O-Q). We found that while WDR90 precisely localizes to the centriolar microtubule wall (Figure 1P, average distance between WDR90 and tubulin: *Δ*= 2nm +/ 12), POC1B, FAM161A, POC5 and Centrin-2 signals were shifted towards the centriole lumen in comparison to the tubulin signal, as previously reported (Figure 1P, *Δ*= 15nm +/- 8; 22nm +/- 5; 27nm +/- 6 and 28nm +/- 9, respectively) (Le Guennec et al., 2020). These results demonstrate that WDR90 longitudinal localization is similar to the inner scaffold components but that WDR90 localizes on the microtubule triplet. This result suggests that WDR90 is a component of the centriolar microtubule triplet of the central core region.

### POC16/WDR90 is an evolutionary conserved microtubule associated protein

Proteins of the POC16/WDR90 family consist of an N-terminal DUF667-containing domain (domain of unknown function), homologous to the ciliary inner junction protein FAP20 (Figure S1A) (Yanagisawa et al., 2014), which is followed by multiple WD40 repeats that form *β*-propeller structures (Figure 2A and Figure S1B) (Xu and Min, 2011).

**Figure 2.**
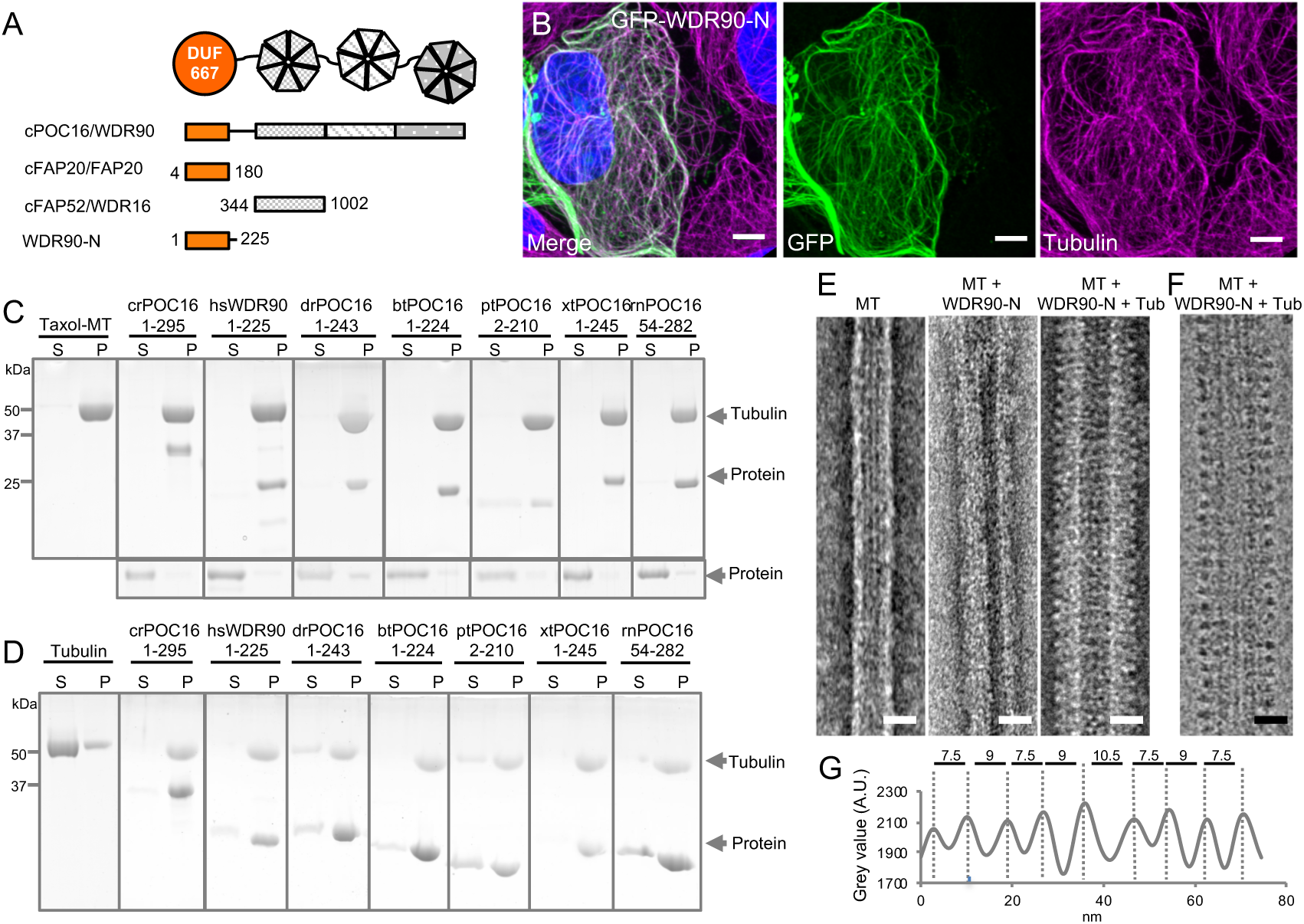
**WDR90/POC16-DUF667 directly binds both microtubules and tubulin. (See also Figures S1-S3).** (A) Schematic of WDR90/POC16 conservation and homologous domains with the *Chlamydomonas* cilia proteins FAP20 and FAP52/WDR16. DUF667 domain is in orange and WD40 repeats are in grey. (B) Human U2OS cells transiently overexpressing GFP-WDR90-N (1-225) stained for GFP (green) and tubulin (magenta). Scale bar: 5µm. (C and D) Coomassie-stained SDS-PAGE of pelleting assays performed *in vitro* with taxol-stabilized microtubules (C), and free tubulin (D), in the presence of different recombinant POC16/WDR90-DUF667 protein orthologs (related to Figure S1A, B). The solubility of proteins alone was assessed in parallel to the microtubule-pelleting assay. All tested proteins were soluble under the tested condition (bottom panel). (E) Electron micrographs of negatively stained taxol-stabilized microtubules alone (MT) or subsequently incubated with recombinant WDR90-N (1-225) alone (MT + WDR90-N) or in combination with tubulin (MT + WDR90-N + Tub). Scale bar: 25nm (F) Cryo-electron micrograph of taxol-stabilized microtubules subsequently incubated with recombinant WDR90-N (1-225) and tubulin. Scale bar: 25nm (G) Periodicity of complexed WDR90-N (1-225)-tubulin oligomers bound to the microtubule shown in (F).

First, we wanted to probe the evolutionary conservation of POC16/WDR90 family members as centriolar proteins. To this end, we raised an antibody against *Paramecium tetraurelia* POC16 and confirmed its localization at centrioles similarly to what we found in *Chlamydomonas reinhardtii* and human cells (Figure S1C) (Hamel et al., 2017).

Further driven by its predicted homology to the microtubule associated protein FAP20 (Khalifa et al., 2019) and the underlying hypothesis that POC16/WDR90 proteins might be joining A and B microtubules as well as by their precise localization on the microtubule wall (Figure 1), we first set out to understand the structural homology between the predicted structures of POC16-DUF667 domain to the recently published near atomic structure of FAP20 from flagella microtubule doublets (Khalifa et al., 2020; Ma et al., 2019) (Figure S2A-C). Strikingly, we observed high similarities between the two structures, suggesting similar biological functions at the inner junction. Moreover, we fitted POC16 model prediction into FAP20 cryo-EM density map and found a good concordance, further hinting for a conserved localization at the level of the microtubule triplet (Figure S2D).

Prompted by this result, we then tested whether POC16/WDR90 proteins, similar to FAP20, can bind microtubules both in human cells as well as *in vitro*. To do so, we overexpressed the N-terminal part of WDR90 and crPOC16 comprising the DUF667 domain (WDR90-N(1-225) and CrPOC16(1-295), respectively) fused to GFP in U2OS cells and found that this region is sufficient to decorate cytoplasmic microtubules (Figures 2B and S3A). We next tested whether overexpressing such a WDR90-N-terminal fragment could stabilize microtubules. To this end, we analyzed the microtubule network in cells overexpressing mCherry-WDR90-N after depolymerizing microtubules through a cold shock treatment (Figure S3B-D). We found that while low expressing cells did not maintain a microtubule network, high expressing cells did. This suggests that WDR90-N can stabilize microtubules. In contrast, we observed that full-length WDR90 fused to GFP only anecdotally binds microtubules. This observation suggests a possible autoinhibition conformation of the full-length protein and/or to interacting partners preventing microtubule binding in the cytoplasm (Figure S3E).

Next, we determined whether different POC16/WDR90 N-terminal domains directly bind to microtubules *in vitro* and whether this function has been conserved in evolution. Bacterially expressed, recombinant POC16/WDR90 DUF667 domains from seven different species were purified and their microtubule interaction ability was assessed using a standard microtubule-pelleting assay (Figure S1A and Figure 2C). We found that every POC16/WDR90 DUF667 domain directly binds to microtubules *in vitro*. This interaction was further confirmed using negative staining electron microscopy, where we could observe recombinant WDR90-N localizing on *in vitro* polymerized microtubules (Figure 2E).

We next investigated whether POC16/WDR90 DUF667 domain could also interact with free tubulin dimers, considering that closure of the inner junction between the A and B microtubules necessitates two microtubule/tubulin-binding sites as recently reported for FAP20 (Ma et al., 2019). We observed that all POC16/WDR90 DUF667 orthologs directly interact with tubulin dimers, generating oligomers that pellet under centrifugation (Figure 2D). We then tested whether the DUF667 domain could still interact with tubulin once bound to microtubules. We subsequently incubated either WDR90-N or crPOC16(1-295) pre-complexed with microtubules with an excess of free tubulin and analyzed their structural organization by electron microscopy (Figure 2E, F and Figure S3F, G). We observed an additional level of decoration due to the simultaneous binding of the DUF667 domains with tubulin and microtubules (Figure 2E, F and Figure S3F, G). Furthermore, we revealed a 8.5 nm periodical organization of tubulin-WDR90-N oligomers on microtubules (Figure 2G), similar to the recent high-resolution structure of the ciliary microtubule doublet showing that monomeric FAP20 interacts with both A- and B-microtubules every 8nm at the inner junction (Khalifa et al., 2020; Ma et al., 2019). In this context, it is tempting to speculate that the DUF667 domain of POC16/WDR90 is also monomeric, however it is also possible that WDR90 forms a homodimer capable of interacting with the microtubules and tubulin.

Based on these results, we concluded that POC16/WDR90 is an evolutionary conserved microtubule/tubulin-interacting protein with the capacity to connect microtubules, a functional prerequisite for an inner junction protein that simultaneously interacts with the A and B microtubules.

### WDR90 is recruited in G2 during centriole core elongation

We next assessed whether WDR90 recruitment at centrioles is correlated with the appearance of inner scaffold proteins during centriole biogenesis. Centrioles duplicate only once per cell cycle during S phase, with the appearance of one procentriole orthogonally to each of the two mother centrioles. Procentrioles then elongate during the following G2 phase of the cell cycle, acquiring the inner scaffold protein POC5 that is critical for the formation of the central and distal parts of the nascent procentriole (Azimzadeh et al., 2009). We followed endogenous WDR90 localization across the cell cycle by analyzing synchronized human RPE1 cells fixed at given time points and stained for either Centrin-2 or HsSAS-6, both early protein marker of duplicating centrioles (Azimzadeh et al., 2009; Strnad et al., 2007) (Figure 3A-F and Figure S4A, B). We found that while Centrin-2 and HsSAS-6 are recruited as expected early on during procentriole formation in S phase, WDR90 starts appearing only in early G2 when procentriole elongation starts (Figure 3A-F). Signal intensity analysis over the cell cycle further demonstrates that WDR90 appears on procentrioles in early G2 and reaches full incorporation by the end of G2 (Figure 3G, H), similarly to the reported incorporation of the inner scaffold protein POC5 (Azimzadeh et al., 2009).

**Figure 3.**
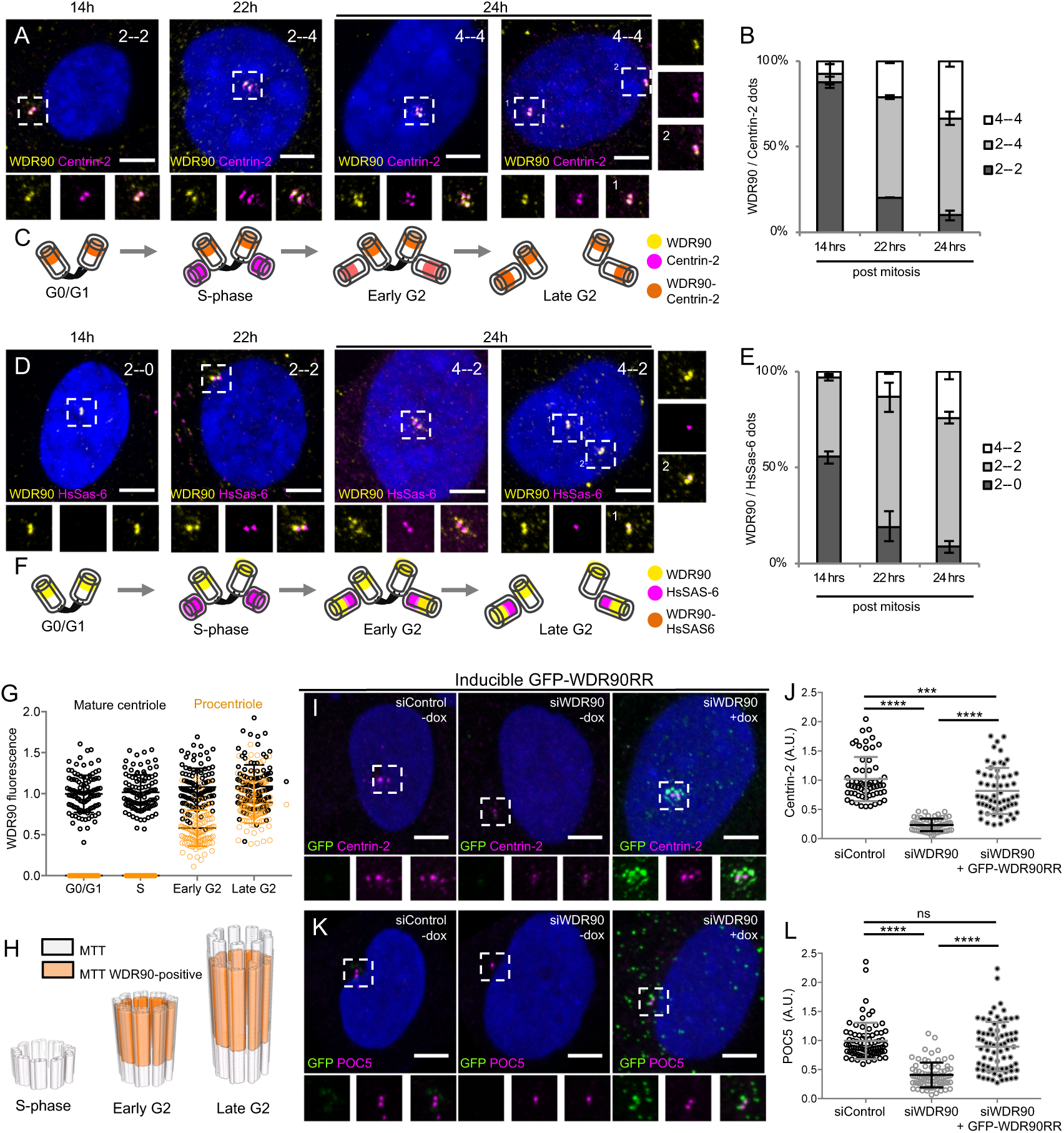
**WDR90 is recruited in G2 and is important for inner scaffold components recruitment to centrioles (See also Figures S4 and S5).** (A) Human RPE1 p53-cells synchronized by mitotic shake-off, fixed at different time points for different cell-cycle stages (related to Fig S4A, B) and stained with WDR90 (yellow) and Centrin-2 (magenta). DNA is in blue. Dotted white squares correspond to insets. Numbers on the top right indicate respectively WDR90 and Centrin-2 numbers of dots. Scale bar: 5µm. (B) Percentage of cells with the following numbers of WDR90/Centrin-2 dots based on A, n=300 cells/condition from 3 independent experiments. Average +/- SD: Refer to **Table 2**. (C) Model for WDR90 and Centrin-2 incorporation during centriole biogenesis based on A. (D) Human RPE1 p53-cells synchronized by mitotic shake-off, fixed at different time points for different cell-cycle stages and stained with WDR90 and HsSAS-6. (E) Percentage of cells with the following numbers of WDR90 and HsSAS-6 based on D, n=300 cells/condition from 3 independent experiments. Average +/- SD: refer to **Table 3**. (F) Model for WDR90 and HsSAS-6 incorporation during centriole biogenesis based on D. (G) WDR90 fluorescence intensity at centrioles according to cell cycle progression, n=45 cells/condition from 3 independent experiments. Black fill represents WDR90 at mature centrioles, orange fill represents WDR90 at procentrioles. (H) Schematic representation of WDR90 incorporation during centriole biogenesis according to cell cycle progression based on G. (I, K) Human U2OS GFP-WDR90 RNAi-resistant version (GFP-WDR90RR) inducible stable cell line treated with control or *wdr90* siRNA and stained for either GFP and Centrin-2 (I) or GFP and POC5 (K) Scale bar: 5µm. Dotted white squares indicate insets. - and + dox indicates induction of GFP-WDR90RR expression. (J) Centrosomal Centrin-2 fluorescence intensity based on I, n= 60 cells/condition from 3 independent experiments. Average +/- SEM (A.U.): Control – dox= 1.02 +/- 0.05, siWDR90 – dox= 0.23+/- 0.01, siWDR90 + dox= 0.82 +/- 0.01. Statistical significance assessed by one-way ANOVA. (L) Centrosomal POC5 fluorescence intensity based on K, n= 75 cells/condition from 3 independent experiments. Average +/- SEM (A.U.): Control – dox= 0.99 +/- 0.04, siWDR90 – dox= 0.41+/- 0.02, siWDR90 + dox= 0.89 +/- 0.05. Statistical significance assessed by one-way ANOVA.

Moreover, we noticed that besides its centriolar distribution, WDR90 localizes also to centriolar satellites, which are macromolecular assemblies of centrosomal proteins scaffolded by the protein PCM1 and involved in centrosomal homeostasis (Drew et al., 2017) (Figure S4C-H). Thus, we tested whether WDR90 satellite localization depends on the satellite protein PCM1 by depleting PCM1 using siRNA and assessing WDR90 distribution. We found that in absence of PCM1, WDR90 is solely found at centrioles (Figure S4E-H), demonstrating that WDR90 satellite localization is PCM1-dependent.

Altogether, these data establish that WDR90 is a centriolar and satellite protein that is recruited to centrioles in the G2-phase of the cell cycle, during procentriole elongation and central core/distal formation, similarly to the recruitment of the inner scaffold protein POC5.

### WDR90 is important to recruit Centrin-2 and POC5

To better understand the function of WDR90, we analyzed cycling human cells depleted for WDR90 using siRNA and co-labeled WDR90 with either the early centriolar marker Centrin-2 or the G2-marker POC5. As previously shown (Hamel et al., 2017), WDR90 siRNA-treated cells showed significantly reduced WDR90 levels at centrosomes in comparison to control cells (Figure S5A, C). Moreover, we observed an asymmetry in signal reduction at centrioles in WDR90-depleted cells, with only one of two Centrin-2 positive centrioles still associated with WDR90 in G1 and early S-phase (69% compared to 10% in controls) and one of four Centrin-2 positive centrioles in S/G2/M cells (77% compared to 0% in controls, Figure S5B). As the four Centrin-2 positive dots indicate duplicated centrioles, this result suggests that the loss of WDR90 does not result from a duplication failure (Figure S5B). We postulate that the remaining WDR90 signal possibly corresponds to the mother centriole and that the daughter has been depleted from WDR90 (Figure S5E), similarly to what has been observed for the protein POC5 (Azimzadeh et al., 2009). We further conclude that WDR90 is stably incorporated into centrioles, in agreement with its possible structural role.

We also noted that the intensity of the Centrin-2 and POC5 signals were markedly reduced upon WDR90 siRNA treatment (Figure S5D-K). Indeed, we found that only 39% of WDR90-depleted cells displayed 2 POC5 dots in G1 (negative for HsSAS-6 signal) in contrast to the 86% of control cells with 2 POC5 dots (Figure S5H). Moreover, 68% of control cells had 2 to 4 POC5 dots in S/G2/M (associated with 2 HsSAS-6 dots) in contrast to 29% in WDR90-depleted condition (Figure S5H). The HsSAS-6 signal was not affected in WDR90-depleted cells, confirming that initiation of the centriole duplication process is not impaired under this condition (Figure S5G, J, L). Similarly, the fluorescence intensity of the distal centriole cap protein CP110 was not changed under WDR90-depletion in contrast to the Centrin-2 signal reduction (Fig S5M-O). These results establish that the localization of Centrin-2 and POC5, two components of the inner scaffold, are affected upon WDR90 depletion in contrast to the proximal protein HsSAS-6 and distal cap protein CP110.

To ascertain this phenotype, we generated a stable cell line expressing a siRNA-resistant version of WDR90 fused to GFP in its N-terminus (GFP-WDR90RR) upon doxycycline induction. We found that expression of GFP-WDR90RR restores partially the Centrin-2 and POC5 signals at centrioles (Figure 3I-L).

Taken together, these results indicate that the depletion of WDR90 leads to a decrease in Centrin-2 and POC5 localization at centrioles but does not affect the initiation of centriole duplication nor the recruitment of the distal cap protein CP110.

### *Chlamydomonas* POC16 is crucial to maintain centriole core integrity

To investigate the structural role of POC16/WDR90 proteins on centrioles, we initially turned to the *Chlamydomonas reinhardtii poc16m504* mutant generated by the Chlamy Library project (Li et al., 2016), which contains an insertion of a CIB1 cassette in the *poc16* genomic sequence (Hamel et al., 2017). We demonstrated previously that this mutant is not a null, but has reduced amount of POC16 proteins detected at centrioles and displays flagella defects with 80% of mutant cells bearing 0, 1 or impaired flagella (Figure S6A-C) (Hamel et al., 2017). We hypothesized from these results that a POC16 truncated protein is made, although this could not be confirmed, as POC16 antibodies do not detect the 230kDa endogenous protein in centriolar extracts by Western Blot (Hamel et al., 2017). Here, we confirmed, by performing immunofluorescence analysis of wild-type and *poc16m504 Chlamydomonas* cells co-stained for POC16 and tubulin, that the overall POC16 levels at centrioles were reduced (Figure 4A, B). Moreover, we noticed that 52% of *poc16m504* centriole pairs had only one detectable POC16 dot and 25% had none as compared to the 2 POC16 dots in the wild-type (Figure 4C). In contrast, by staining for the cartwheel component Bld12 (Nakazawa et al., 2007), we found that the fluorescent signal was similar to wild-type in this background, suggesting that the proximal region of the centriole is not affected (Figure S6D-F), similarly to human cells.

**Figure 4.**
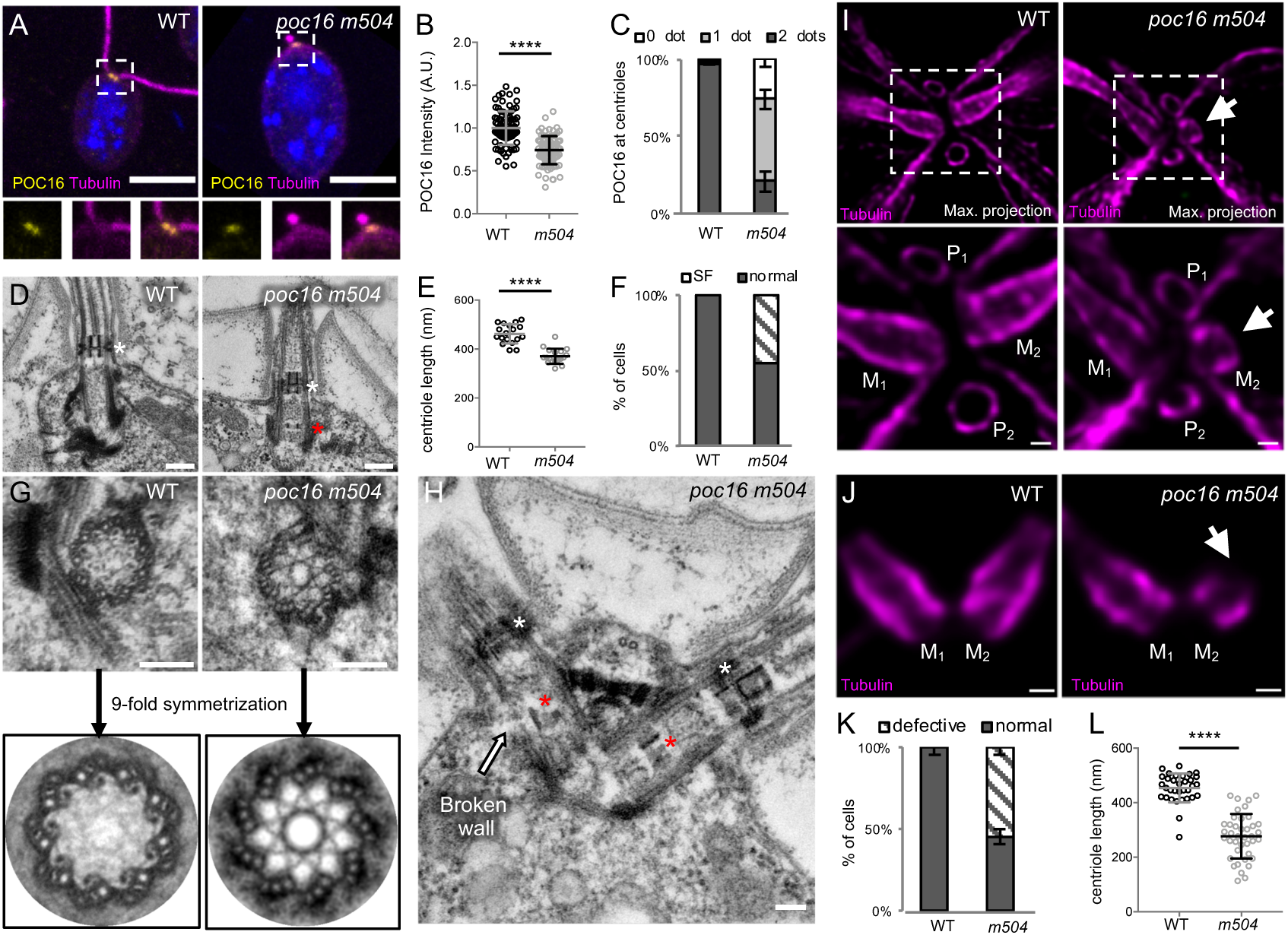
**POC16 mutant lacks the inner scaffold (See also Figure S6).** (A) Confocal image of *Chlamydomonas* wild-type (WT) and *poc16 m504* mutant stained for tubulin (magenta) and POC16 (yellow), DNA is in blue. Dotted squares correspond to insets. Scale bar: 5µm. (B) POC16 fluorescence intensity based on A, n=90 cells/condition from 3 independent experiments. Average +/- SD: WT: 1+/- 0.2 (A.U.), *poc16m504*: 0.74 +/- 0.2 (A.U.) Normality assessed by Pearson test, Welch T test p< 0.0001. (C) Percentage of cells displaying two, one or no POC16 dots per cell, n=300 cells/condition from 3 independent experiments. Average +/- SD: WT 2 dots: 97% +/- 0.5, 1 dot: 1.3% +/- 1.5, no dot: 1.3% +/- 1.4; *poc16 m504* 2 dots: 20.1% +/- 6.4, 1 dot: 52% +/- 6.1, no dot: 25.2% +/- 4.5. (D) Electron micrograph of *Chlamydomonas* WT and *poc16 m504* sections revealing the presence of ectopic stellate fibers (SF, white star: normal position in the transition zone; red star: ectopic localization of SF within the central core region of centrioles) inside the lumen of *poc16 m504* centrioles. Scale bar: 250nm (E) Centriole length in WT and *poc16m504* cells, 18 centrioles analyzed in each condition. Average +/- SD: WT= 462 +/- 9nm, *poc16m504*= 371 +/- 7nm. Normality assessed by Pearson test, Welch T test p< 0.0001. (F) Percentage of centrioles with ectopic stellate fibers (SF) in WT (0%) and *poc16 m504* (46%), 18 centrioles analyzed in each condition. (G) Electron micrographs of transversal section of *Chlamydomonas* WT (left) and *poc16 m504* (right) centrioles (top panel) and their corresponding circularized and symmetrized version (bottom panel). Top views circularization and symmetrization were performed using CentrioleJ. Scale bar: 200nm (H) *poc16 m504* mutant displaying a broken centriolar microtubule wall (white arrow). Note the SF in the transition zone (white star) as well as the ectopic SF (red star) within the central core region of *poc16m504* centrioles. Scale bar: 200nm (I) *In cellulo Chlamydomonas* WT or *poc16m504* centrioles/flagella expanded using U-ExM and stained for tubulin (magenta). M stands for mature centriole and P for procentriole. Arrows point to defective mature centrioles. Scale bar: 100nm (J) *In cellulo Chlamydomonas* WT or *poc16m504* pair of mature centrioles expanded using U-ExM and stained for tubulin (magenta). Arrows point to defective mature centrioles. Scale bar: 100nm (K) Percentage of cells with abnormal mature centrioles. Average +/- SD: WT 0% +/- 0, *poc16m504*: 55% +/- 5 from 3 independent experiments. (L) Centriolar length based on J, n= 30 centrioles/condition from 3 independent experiments. Average +/- SD: WT 454nm +/- 53, *poc16m504*: 277nm +/- 82. Mann-Whitney test p< 0.0001.

To assess whether the ultrastructure and in particular the central core region of centrioles in *poc16m504* cells was defective, we analyzed this mutant using electron microscopy of resin-embedded specimens (Figure 4D-H). We first noticed that the *poc16m504* mutant displayed shorter centrioles with an average length of 370 nm (+/- 7 nm) compared to 460 nm (+/- 9 nm) in the wild type (Figure 4D, E). Moreover, we found that the stellate fibers present in the transition zone of wild-type centrioles (Figure 4D, white star) (Geimer and Melkonian, 2004), are ectopically localized to the central core region of *poc16m504* mutants in 46% of the cases (Figure 4D, F-H, red star). This additional localization of stellate fibers has previously been described for the *δ*-tubulin mutant *uni-*3, which also displays defective microtubule triplets (O’Toole et al., 2003). However, in contrast to *uni-3*, we noted that microtubule triplets were apparently not affected in the *poc16m504* mutant (Figure 4G). However, even if extremely rare owing to the difficulty to obtain a perfect longitudinal view of the centriole in the resin-embedded sections, we observed a centriole with a broken microtubule wall at the level of the central core region, suggestive of centriole fracture (Figure 4H, arrow).

To better characterize this phenotype, we turned to U-ExM that allows visualization of centrioles ultrastructure in a more quantitative manner and in the context of the whole organism (Gambarotto et al., 2019). While the procentrioles looked intact, confirming that proximal assembly initiation is not affected in this mutant, 55% of the *poc16m504* mutants displayed defective mature centrioles compared to wild type (Figure 4I-K and Figure S6G, H). We notably observed incomplete mature centrioles lacking the entire central and distal parts, suggesting either a defect in centriole assembly and elongation or an impairment of centriole stability (Figure 4I, J and Figure S6G, H). Moreover, consistent with our electron microscopy analysis, quantification of mature centrioles in this mutant demonstrated that centrioles are shorter (Figure 4L).

Altogether, these results demonstrate that the central core region of *poc16m504* mutants is impaired, highlighting a potential role of POC16 in either maintaining the structural integrity of the microtubule wall in this region of the centriole and/or its assembly.

### WDR90 depletion leads to a loss of inner scaffold components and to centriole fracture

Based on these findings, we wondered whether WDR90 depletion might lead to a loss of inner scaffold components as well as to a centriole architecture destabilization in human cells. We tested this hypothesis by analyzing centrioles from WDR90-depleted U2OS cells using U-ExM (Figure 5). As expected, we observed a strong reduction of WDR90 at centrioles, with a reminiscent asymmetrical signal in one of the two mature centrioles (Figure 5A-C). Unexpectedly, we found that WDR90-depleted centrioles exhibited a slight tubulin length increase (502 nm +/- 65 compared to 434 nm +/- 58 in controls), potentially indicative of a defect in centriole length regulation (Figure 5D). In contrast, despite a slight decrease at the level of the central core, we did not observe, in neither of the conditions, any significant difference in centriole diameter at the proximal and very distal regions (Figure 5E).

**Figure 5.**
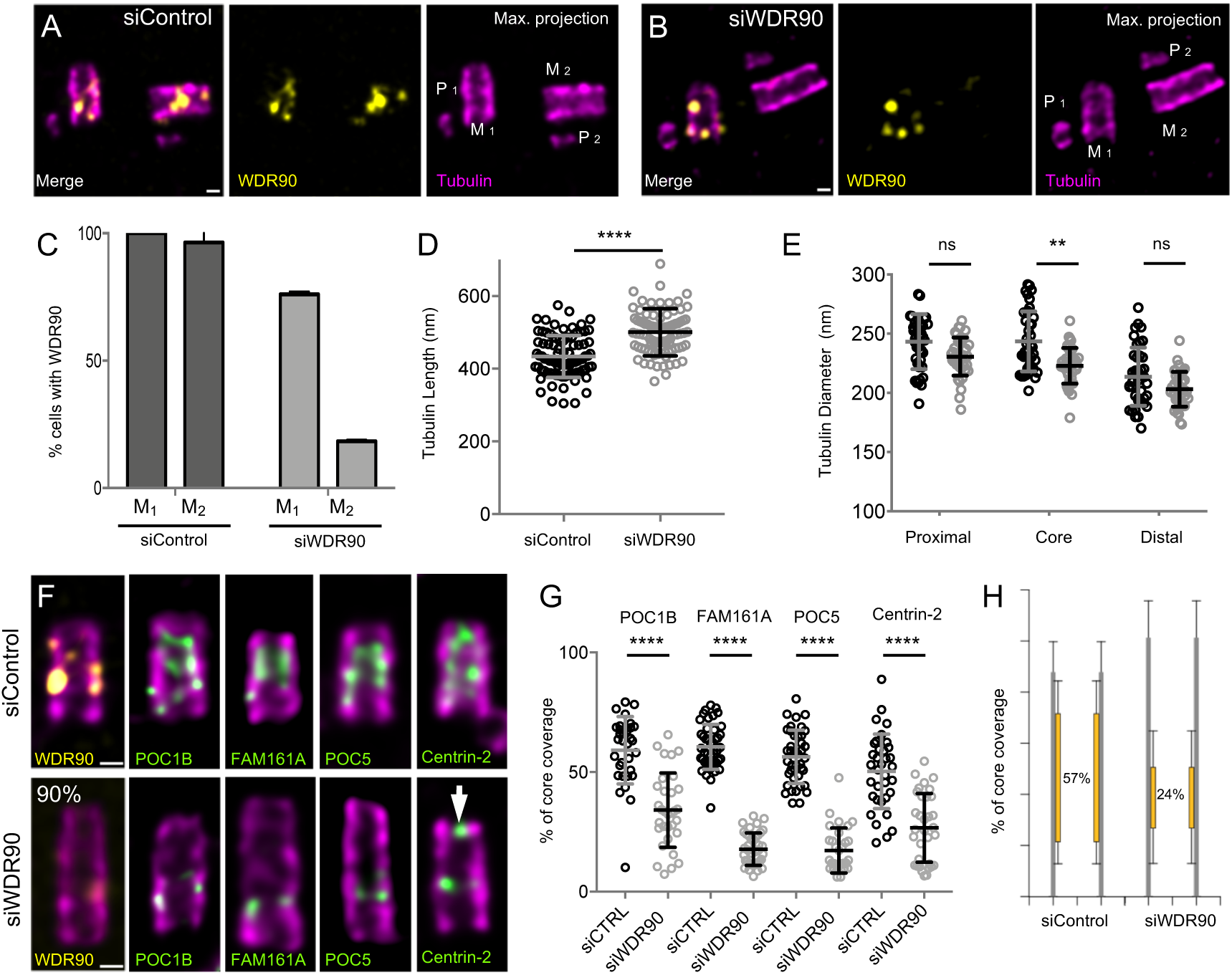
**WDR90 is crucial for inner scaffold components localization (see also Figure S7).** (A, B) *In cellulo* U-ExM expanded centrioles from S-phase U2OS treated with either scramble (A) or *wdr90* siRNA (B) stained for tubulin (magenta) and WDR90 (yellow). M stands for mature centriole and P for procentriole. Scale bar: 100nm. (C) Percentage of cells with WDR90 at mature centrioles, n=90 cells/condition from 3 independent experiments Average +/- SD: siControl= M1: 100% +/- 0, M2: 96% +/- 4.7, siWDR90= M1: 76% +/- 1, M2: 18% +/- 0.6. (D) Tubulin length in nm, n=90 centrioles/condition from 3 independent experiments. Average +/- SD: siControl= 434nm +/- 58, siWDR90= 500nm +/- 65. Mann-Whitney p<0.0001. (E) Tubulin diameter measured in the proximal, central core and distal regions of expanded centrioles in siControl (black circles) and *wdr90* siRNA (siWDR90, grey circles). n= 42 and 43 centrioles for siControl and siWDR90 respectively from 2 independent experiments. Averages +/- SD: refer to **Table 4**. Statistical significance assessed by one-way ANOVA. (F) *In cellulo* U-ExM expanded U2OS centrioles treated with either scramble or *wdr90* siRNA stained for tubulin (magenta) and WDR90 (yellow) or POC1B, FAM161A, POC5 or Centrin-2 (inner scaffold components: green). Scale bar: 100nm. (G) Inner scaffold protein length, n>30 centrioles/condition from 3 independent experiments. Average +/- SD: refer to **Table 5**. Statistical significance assessed by one-way ANOVA. (H) Average core length coverage. Average +/- SD: siControl= 57% +/- 13; siWDR90= 24% +/- 14.

We next analyzed whether the localization of the four described inner scaffold components POC1B, FAM161A, POC5 and Centrin-2 would be affected in WDR90-depleted cells. We found that the localization of these four proteins in the central core region of centrioles was markedly altered in WDR90-depleted centrioles (Figure 5F, G). Instead of covering ∼60% of the entire centriolar lumen, we only observed a ∼20% remaining belt, positive for inner scaffold components at the proximal extremity of the core region (Figure 5F-H and Figure S7A, B), suggesting that their initial recruitment may not be entirely affected. Another possibility would be that incomplete depletion of WDR90 allows for partial localization of inner scaffold components. It should also be noted that Centrin-2, which displays a central core and an additional distal tip decoration (Le Guennec et al., 2020), was affected specifically in its inner core distribution (Figure 5F, white arrow, Figure S7A, B).

The discovery of the inner scaffold within the centriole led to the hypothesis that this structure is important for microtubule triplet stability and thus overall centriole integrity (Le Guennec et al., 2020). In line with this hypothesis, we found that upon WDR90 depletion, 10% of cells had their centriolar microtubule wall broken, indicative of microtubule triplets fracture and loss of centriole integrity (15 out of 150 centrioles, Figure 6, Videos 1 and 2). The break occurred mainly above the remaining belt of inner scaffold components (Figure 6A-D), possibly reflecting a weakened microtubule wall in the central and distal region of the centriole. We also noticed that the perfect cylindrical shape (defined as roundness) of the centriolar microtubule wall was affected with clear ovoid-shaped or opened centrioles seen from near-perfect top view oriented centrioles (Figure 6E, F and Figure S7C), illustrating that loss of WDR90 and the inner scaffold leads to disturbance of the characteristic centriolar architecture.

**Figure 6.**
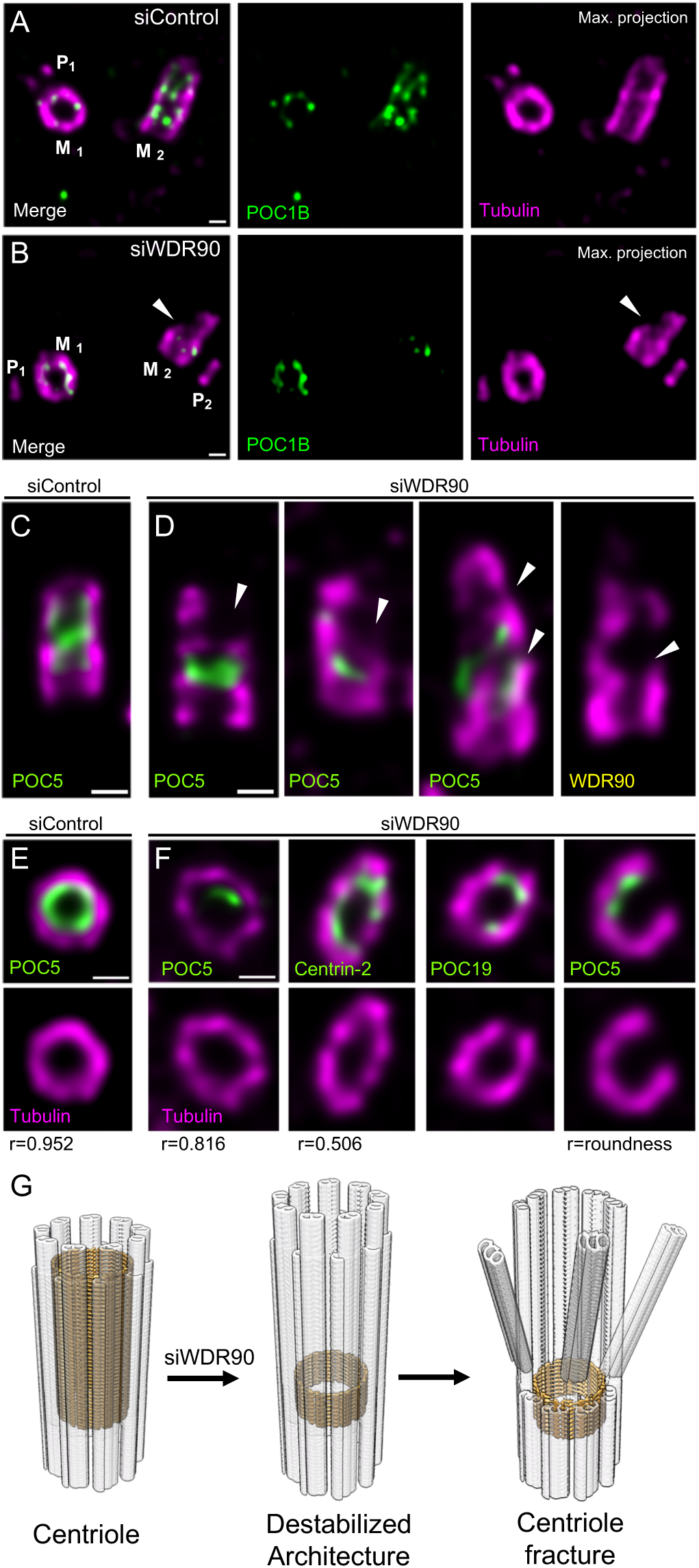
**WDR90 is important for centriole architecture integrity (see also Figure S7, Videos 1 and 2).** (A, B) *In cellulo* U-ExM expanded centrioles from S-phase U2OS cells treated with scramble (A) or *wdr90* siRNA (B), stained for tubulin (magenta) and POC1B (green). White arrowhead: broken microtubule wall of the mature centriole. P: procentriole, M: mature centriole. Scale bars: 100nm. (C, D) *In cellulo* U-ExM expanded centrioles from U2OS cells treated with scramble (C) or *wdr90* siRNA (D), stained for tubulin (magenta) and inner scaffold proteins (green, the specific inner core proteins used for each example is written in each panel), displaying microtubule wall fractures (white arrowheads), lateral view. (E, F) Top views of U-ExM expanded centrioles from U2OS cells treated with scramble. Scale bars: 100nm. (E) or *wdr90* siRNA (F) stained as specified above. Note the loss of roundness of centrioles treated with *wdr90* siRNA. (G) Proposed model of WDR90 function holding microtubule triplets in the central core region of centrioles.

Collectively, these data demonstrate that WDR90 is crucial to ensure inner core protein localization within the centriole core, as well as maintaining the microtubule wall integrity and the overall centriole roundness and stability (Figure 6G).

## Discussion

What maintains centriole barrel stability and roundness is a fundamental open question. Centrioles are microtubule barrel structures held together by the A-C linker at their proximal region and a recently discovered inner scaffold in the central/distal region (Le Guennec et al., 2020). The presence of such an extended scaffold covering 70% of the centriolar length has led to the hypothesis that this structure is important for maintaining centriole integrity (Le Guennec et al., 2020). Our work demonstrates that POC16/WDR90 family proteins, which are important for cilia and flagella formation, constitute an evolutionary conserved central core microtubule triplet component that is essential for maintaining the inner centriolar scaffold proteins. The depletion of WDR90 leads to centriolar defects and impairment of microtubule triplets organization resulting in the loss of the canonical circular shape of centrioles. We also found that this overall destabilization of the centriole can lead to microtubule triplet breakage. Whether this phenotype arises as a consequence of the loss of the inner scaffold or due to the destabilization of the inner junction of the microtubule triplet is still an opened question that should be addressed in the future.

We demonstrate that POC16/WDR90 is a component of the microtubule triplet restricted to the central core region. In addition and based on the sequence and structural similarity to the DUF667 domain of FAP20 that composes the inner junction in flagella, we propose that POC16/WDR90 localizes at the inner junction of the A and B microtubule of the centriolar microtubule triplet. The fact that WDR90 localization is restricted to the central core region led us to hypothesize that another protein, possibly FAP20 as it has been previously reported at centrioles (H. Yanagisawa et al., 2014), could mediate the inner junction between A- and B-microtubule in the proximal region of the centriole. Moreover, in POC16/WDR90 proteins, the DUF667 domain is followed by a WD40 domain sharing a homology with the flagellar inner B-microtubule protein FAP52/WDR16 (Owa et al., 2019) leading us to postulate that the WD40 domains of POC16/WDR90 might also be located inside the B-microtubule of the triplet. However, whether this is the case remains to be addressed in future studies.

Our work further establishes that WDR90 is recruited to centrioles in G2 phase of the cell cycle concomitant with centriole elongation and inner central core assembly. We found that WDR90 depletion does not impair centriole duplication nor microtubule wall assembly, as noted by the presence of the proximal marker HsSAS-6 and the distal cap CP110. In stark contrast, WDR90 depletion leads to a strong reduction of inner scaffold components at centrioles, as well as some centriole destabilization.

Although several examples of centriole integrity loss have been demonstrated in the past, the molecular mechanisms of centriole disruption are not understood. For instance, Delta- and Epsilon-tubulin null human mutant cells were shown to lack microtubule triplets and have thus unstable centrioles that do not persist to the next cell cycle (Wang et al., 2017). Remarkably, these centrioles can elongate with a proper recruitment of the cartwheel component HsSAS-6 and the distal marker CP110 but fails to recruit POC5, a result that is similar to our findings with WDR90 depleted cells. As Delta- and Epsilon-tubulin null human mutant cells can solely assemble microtubule singlets (Wang et al., 2017), we speculate that WDR90 might not be recruited in these centrioles, as the A-and B-microtubule inner junction would be missing. As a consequence, the inner scaffold proteins will not be recruited, as already shown for POC5, leading to the observed futile cycle of centriole formation and disintegration (Wang et al., 2017). It would therefore be interesting to study the presence of WDR90 in these null mutants as well as the other components of the inner scaffold in the future.

Our work also showed that POC16 and WDR90 depletion affects centriole length both in *Chlamydomonas reinhardtii* and human cells. While we observed shorter centrioles in *poc16m504* mutants and the opposite, longer centrioles, in WDR90-depleted cells, these results emphasize the role of POC16/WDR90 in overall centriole length regulation and suggest an unexpected role of the inner scaffold structure in centriole length control. The observed discrepancy between the two phenotypes could arise from species differences or from the fact that we analyzed a mutant (truncated protein) in the case of *Chlamydomonas reinhardtii* POC16 versus an RNAi-mediated depletion in the case of human WDR90. Regardless, it would be of great interest to understand if and how the absence of the inner scaffold can affect the length of the centriole without affecting distal markers such as CP110, which remains unchanged in our experiments. It is very likely that the concomitant elongation of the centriole with the appearance of inner scaffold components in G2 can act on the final length of this organelle.

Given the importance of centriole integrity in enabling the proper execution of several diverse cellular processes, our work provides new fundamental insights into the architecture of the centriole, establishing a structural basis for centriole stability and the severe phenotypes that arise when lost.

## Acknowledgments

We thank Michel Bornens, Eloise Lequoy-Bertiaux and Nikolai Klena for critical reading of the manuscript. We thank Dr. Kanh Huy Bui for sharing his cryo-EM map EMD-20858. We thank the BioImaging Center and PFMU at Unige. We thank Juliette Azimzadeh for sharing the construct GFP-WDR90-FL. We thank the PhD Booster Team of the University of Geneva attributed to E.S. This work is supported by the Swiss National Science Foundation (SNSF) PP00P3_157517 (to P.G.) and 31003A_166608 (to M.O.S.), and by the European research Council ERC ACCENT StG 715289 attributed to Paul Guichard.

## Author contributions

E.S. performed and analyzed all the experiments of the paper except for Figure 2C-D. V.H. and Pa.G. conceived, supervised and designed the project. M.O.S. supervised the biochemical microtubule/tubulin-binding experiments. V.H., Pa.G. and E.S. wrote the manuscript with the input from all authors. D.G. and M.L. contributed to U-ExM experiments. C.Z., and N.O. expressed and purified the recombinant proteins used in this study, performed the experiments presented in Figure 2C-D and generated Figure S1A, B. V.O. (together with N. O.) worked on the POC16 model prediction (Figure S2). S.B. provided technical support for the entire study. M.L.G. provided cryo-EM maps and helped with U-ExM data analysis. F.K. and A-M.T. provided expertise and help for the work performed in *Paramecium tetraurelia* (Figure S1C).

## Supplementary Figure legends

**Figure S1.**
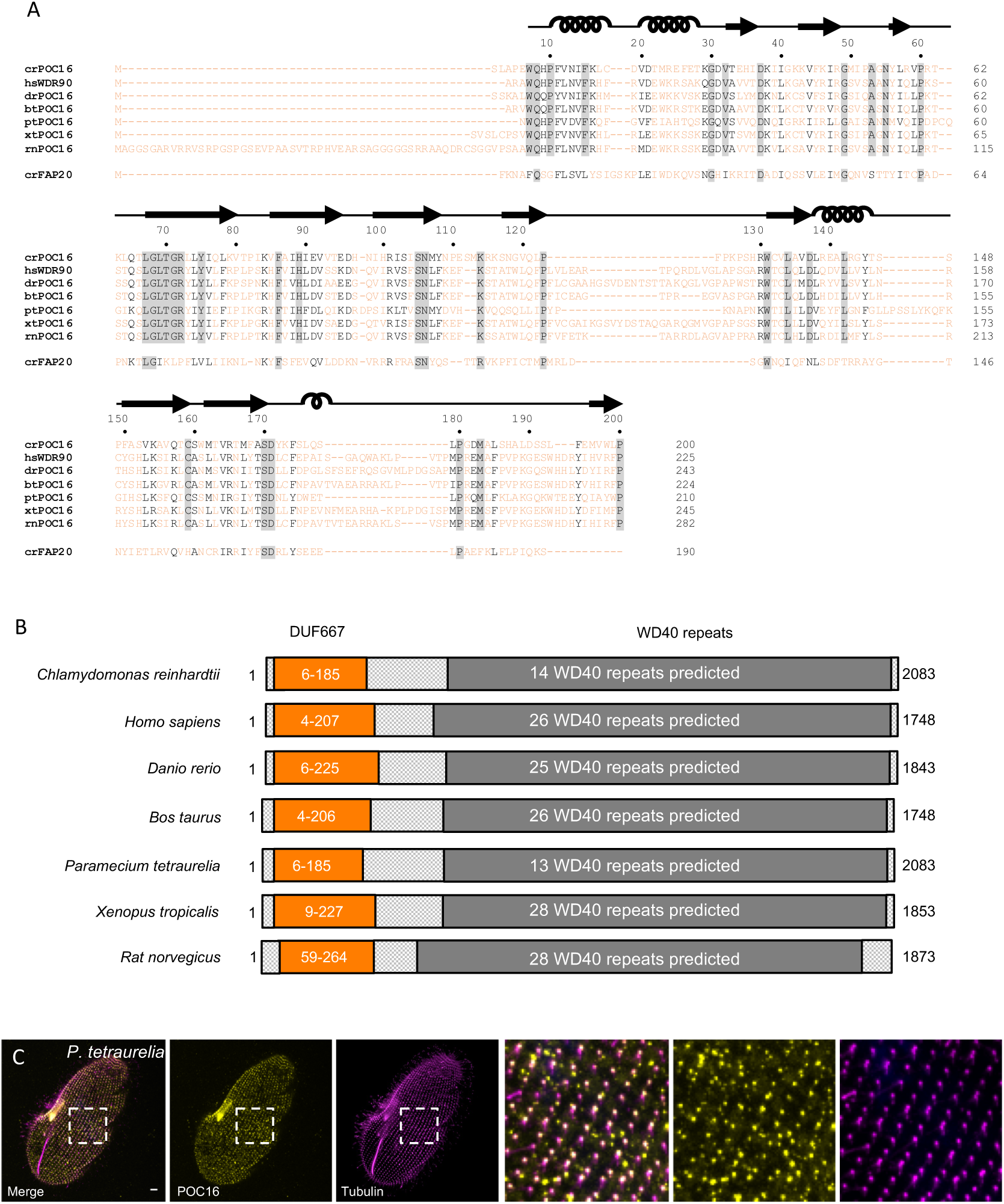
**POC16 conservation across species.** (A) POC16 orthologs DUF667 domain amino acids sequence alignment from 7 different species: *Chlamydomonas reinhardtii* crPOC16(1-200); *homo sapiens* hsWDR90(1-225), *Danio rerio* drPOC16(1-243), *Bovine taurus* btPOC16(1-224), *Paramecium tetraurelia* ptPOC16(1-210), *Xenopus tropicalis* xtPOC16(1-245) and *Rat norvegicus* rtPOC16(1-282). Note also below the alignment with *Chlamydomonas reinhardtii* crFAP20. The secondary structures *α*-helices and *β*-strand are indicated on top of the amino acid sequences. (B) POC16 orthologs domain mapping and conservation. Orange: DUF667 domain. Dark grey: WD40 repeats. (C) *Paramecium tetraurelia* cell fixed and stained for ptPOC16 (yellow) and tubulin (1D5) (magenta), showing that ptPOC16 is a centriolar component. Scale bare: 10µm.

**Figure S2.**
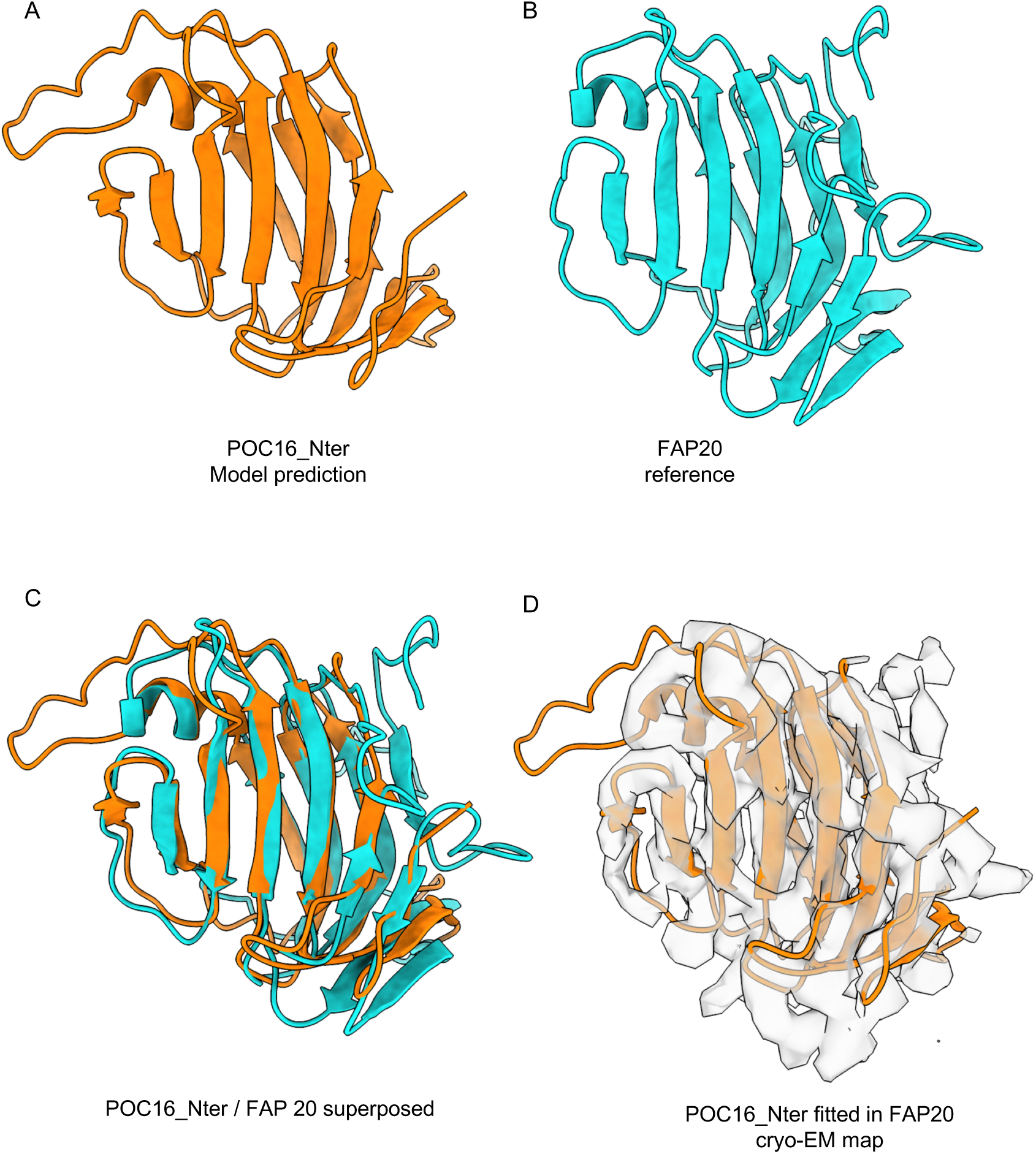
**Model prediction of POC16 Nter.** (A) POC16 3D model and (B) FAP20 reference structure model (Khalifa et al., 2019). (C) Fitting of POC16 against FAP20 yielding a RMSD value of 1.6 Angs. (D) Fitting of the POC16 model excluding the flexible loops in the FAP20 cryo-EM electron density map.

**Figure S3.**
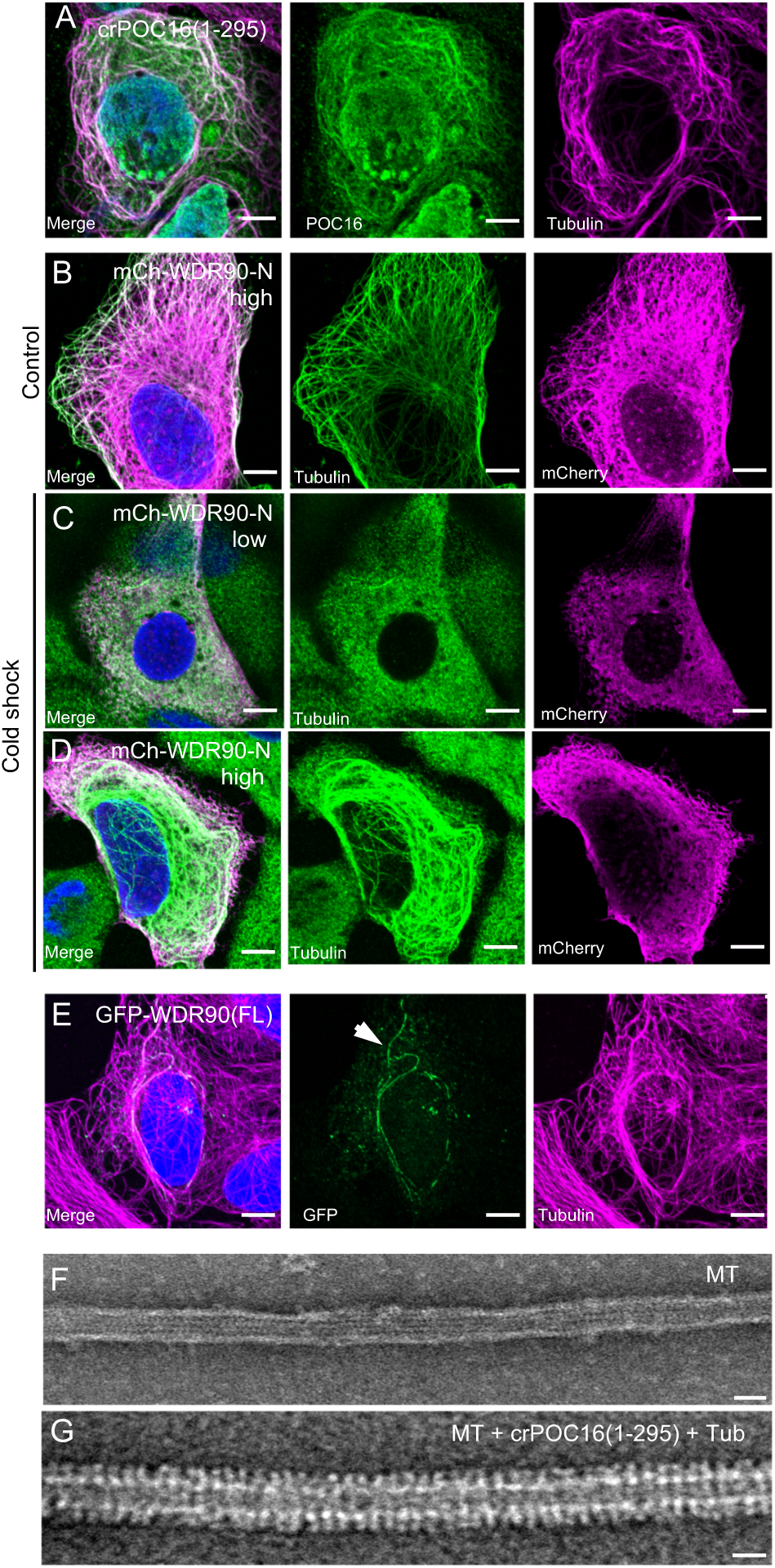
**POC16 and WDR90 bind microtubules.** (A) Human U2OS cells transiently overexpressing GFP-crPOC16(1-295) stained for POC16 (green) and tubulin (magenta). Scale bars for panels A-E: 5µm. (B) Human U2OS cells transiently overexpressing at high level mCherry-WDR90-N(1-225), fixed in control condition and stained for tubulin (green) and mCherry (magenta). (C) Human U2OS cells transiently overexpressing at low level mCherry-WDR90-N(1-225), fixed after 1h of cold shock treatment and stained for tubulin (green) and mCherry (magenta). (D) Human U2OS cells transiently overexpressing at high level mCherry-WDR90-N(1-225), fixed after 1h of cold shock treatment and stained for tubulin (green) and mCherry (magenta). (E) Human U2OS cells transiently overexpressing GFP-WDR90(FL) stained for GFP (green) and tubulin (magenta). Arrowhead indicates WDR90-decorated microtubules. (F) Electron micrograph of negatively stained *in vitro* taxol-stabilized microtubules. Scale bar: 25nm (G) Electron micrograph of negatively stained *in vitro* taxol-stabilized microtubules incubated with recombinant POC16(1-295) and free tubulin. Scale bar: 25nm.

**Figure S4.**
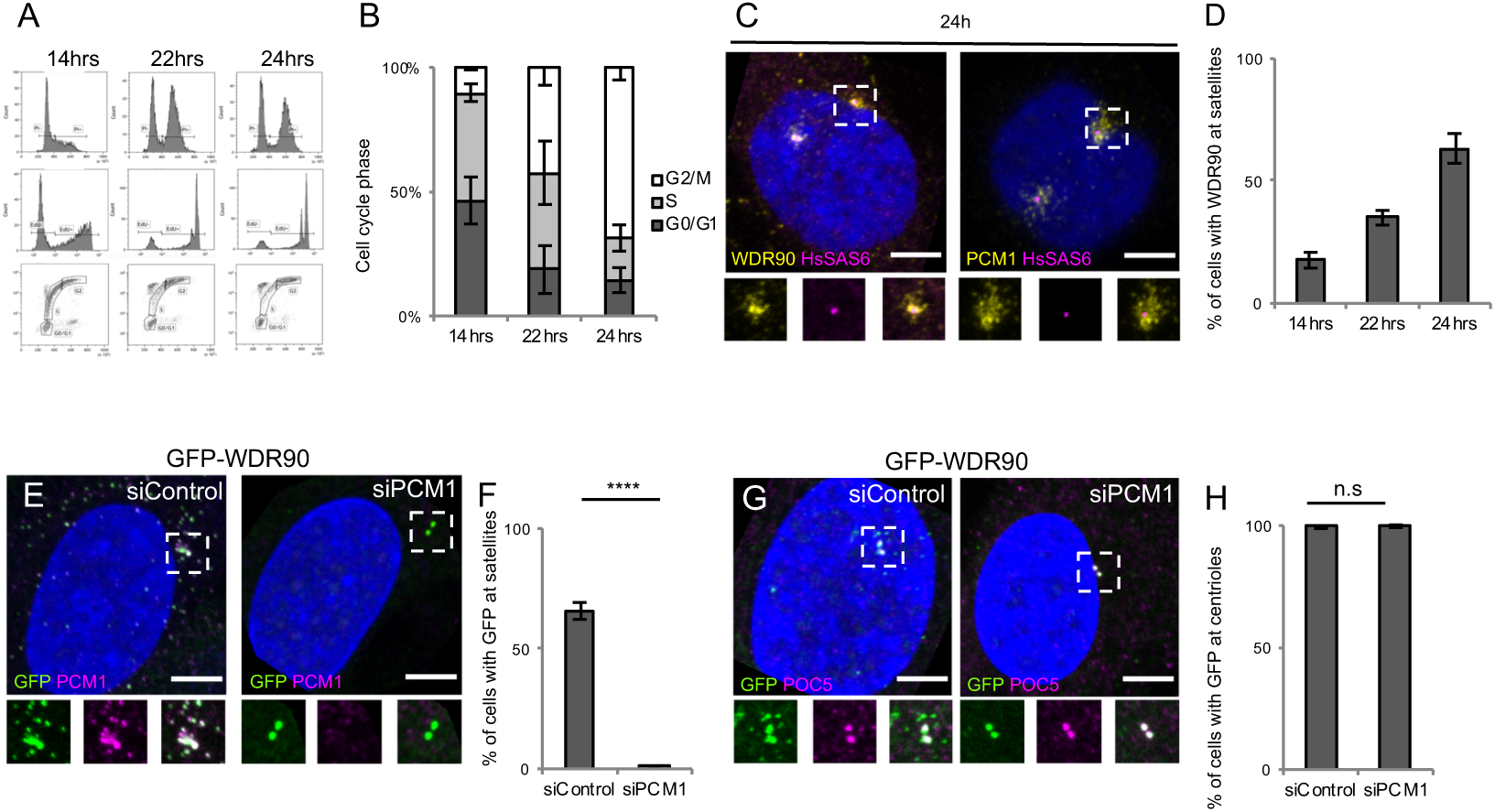
**WDR90 is a satellite and centriolar protein.** (A) FACS profiles of RPE1 p53-cells at different time point post mitotic shake-off, plotted based on propidium iodide (PI) and 5-ethynyl-2′-deoxyuridine (EdU) content. Related to Figure 5(A-E). (B) Percentage of cells in G0/G1, S or G2/M phase based on A, n=25000 cells/condition from 3 independent experiments. Average +/- SD: Refers to **Table 6**. (C) Human RPE1 p53-fixed 24 hours post mitosis and stained for WDR90 (yellow) and HsSAS-6 (magenta) or PCM1 (yellow) and HsSAS-6 (magenta). DNA is in blue. Scale bar: 5µm. Dotted white squares indicate insets. (D) Percentage of cells displaying WDR90 satellite pattern based on C, n=150 cells/condition from 3 independent experiments. Average +/- SD: 14hrs: 18% +/- 3, 22hrs: 35% +/- 3, 24hrs: 63% +/- 6. (E) Human U2OS cells expressing GFP-WDR90 treated with scramble or *pcm1* siRNA and stained for GFP and PCM1. Scale bar: 5µm. Dotted white squares indicate insets (F) Percentage of cells with GFP-WDR90 at satellites based on F, n=300 cells/condition from 3 independent experiments Average +/- SD: siControl: 66% +/- 4, siPCM1: 1% +/- 1. Welch T-test p<0.0001. (G) Human U2OS cells expressing GFP-WDR90 treated with scramble or *pcm1* siRNA and stained for GFP and POC5. Scale bar: 5µm. Dotted white squares indicate insets. (H) Percentage of cells with GFP-WDR90 at centrioles based on H, n=300 cells/condition from 3 independent experiments Average +/- SD: siControl: 99% +/- 1, siPCM1: 100% +/- 1. Welch T-test p=0.5185.

**Figure S5.**
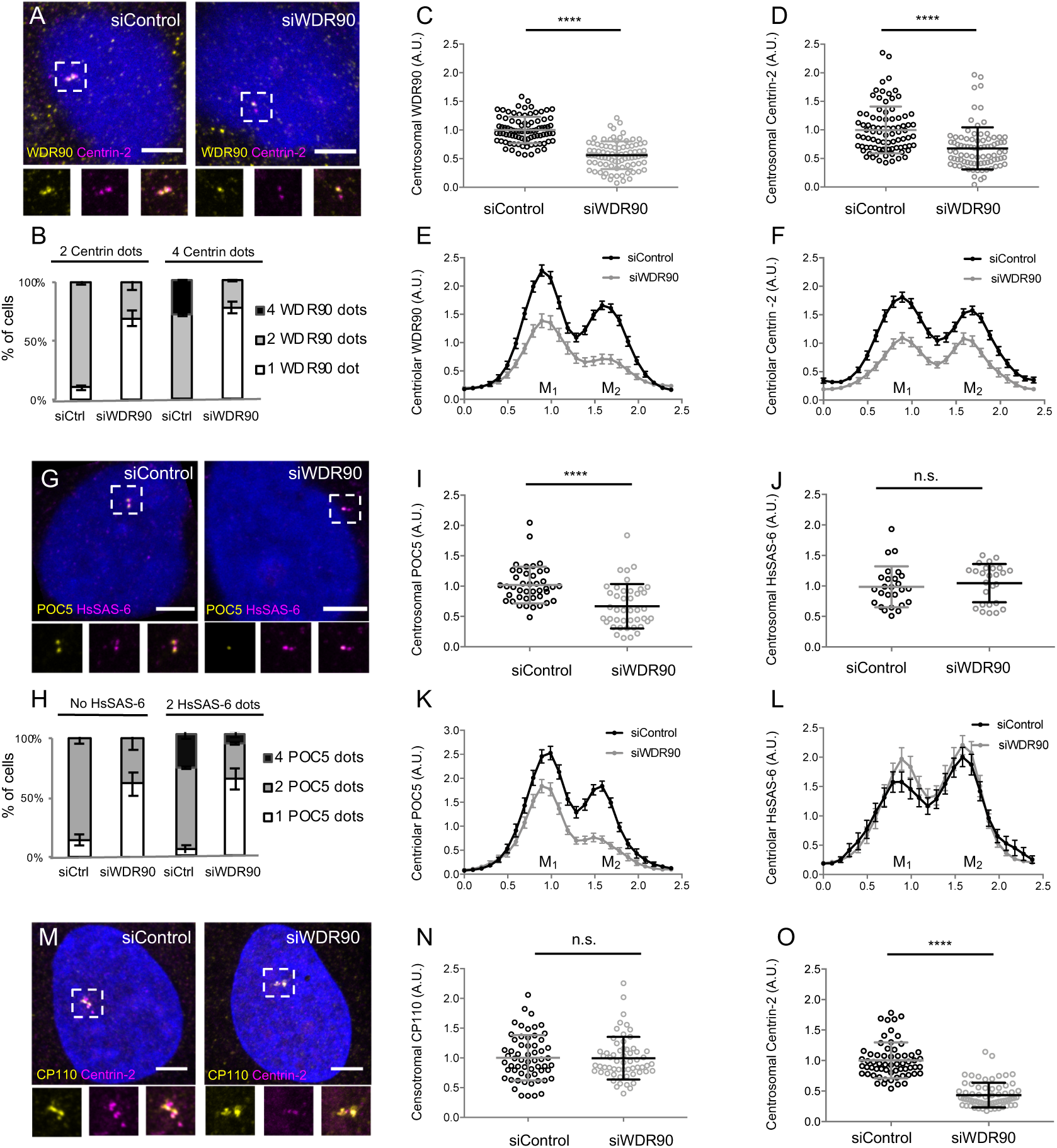
**Depletion of WDR90 impairs Centrin-2 and POC5 localization at centrioles.** (A) Human U2OS cell treated with either scramble or *wdr90* siRNA and stained for WDR90 (yellow) and Centrin-2 (magenta). DNA is in blue. Dotted white squares indicate insets. Scale bar: 5µm. (B) Percentage of cells with the following number of WDR90 dots according to the number of Centrin-2 dots per cell based on A, n=150 cells/condition from 3 independent experiments. Average +/- SD: Refer to **Table 7** (C) WDR90 centrosomal intensity based on A, n=90 cells/condition from 3 independent experiments. Average +/- SD: siControl: 1 +/- 0.2 (A.U.), siWDR90: 0.56 +/- 0.2 (A.U.). Normality assessed by Pearson test, Welch T-test p<0.001. (D) Centrin-2 centrosomal intensity based on A, n=90 cells/condition from 3 independent experiments. Average +/- SD: siControl 1 +/- 0.4 (A.U.), siWDR90: 0.68 +/- 0.4 (A.U). Mann-Whitney p<0.001. (E) Plot profiles of WDR90 centriolar intensity based on A, n=90 cell/condition from 3 independent experiments. M_1_ and M_2_ respectively refer to each mature centriole within pairs (F) Plot profiles of Centrin-2 centriolar intensity based on A, n=90 cells/condition from 3 independent experiments. (G) Human U2OS cell treated with either scramble or *wdr90* and stained for POC5 (yellow) and HsSAS6 (magenta). DNA is in blue. Dotted white squares indicate insets. Scale bar: 5µm. (H) Percentage of cells with the following numbers of POC5 dots according to the number of HsSAS-6 dots per cell based on G, n=150 cells/condition from 3 independent experiments. Average +/- SD: Refer to **Table 8** (I) POC5 centrosomal intensity based on G, n=45 cells/condition from 3 independent experiments. Average +/- SD: siControl 1 +/- 0.3(A.U.), siWDR90: 0.67 +/- 0.4(A.U). Mann-Whitney p<0.001. (J) HsSAS-6 centrosomal intensity based on G, n=30 cells/condition from 3 independent experiments. Average +/- SD: siControl 0.99 +/- 0.3 (A.U.), siWDR90: 1 +/- 0.3 (A.U). Mann-Whitney p=0.2551. (K) Plot profiles of POC5 centriolar intensity based on G, n=45 cells/condition from 3 independent experiments. (L) Plot profiles of HsSAS-6 centriolar intensity based on G, n=30 cells/condition from 3 independent experiments. (M) Human U2OS cell treated with either scramble or *wdr90* siRNA and stained for CP110 (yellow) and Centrin-2 (magenta). DNA is in blue. Dotted white squares indicate insets. Scale bar: 5µm. (N) CP110 centrosomal intensity based on M, n=60 cells/condition from 3 independent experiments. Average +/- SD: siControl 1 +/- 0.4 (A.U.), siWDR90: 0.99 +/- 0.4 (A.U). Mann-Whitney p=0.7756. (O) Centrin-2 centrosomal intensity based on M, n=55 cells/condition from 3 independent experiments. Average +/- SD: siControl 1 +/- 0.3 (A.U.), siWDR90: 0.4 +/- 0.2 (A.U). Mann-Whitney p<0.0001. Note that Centrin-2 intensity served as an internal control for the efficient depletion of WDR90 by siRNA in this experiment.

**Figure S6.**
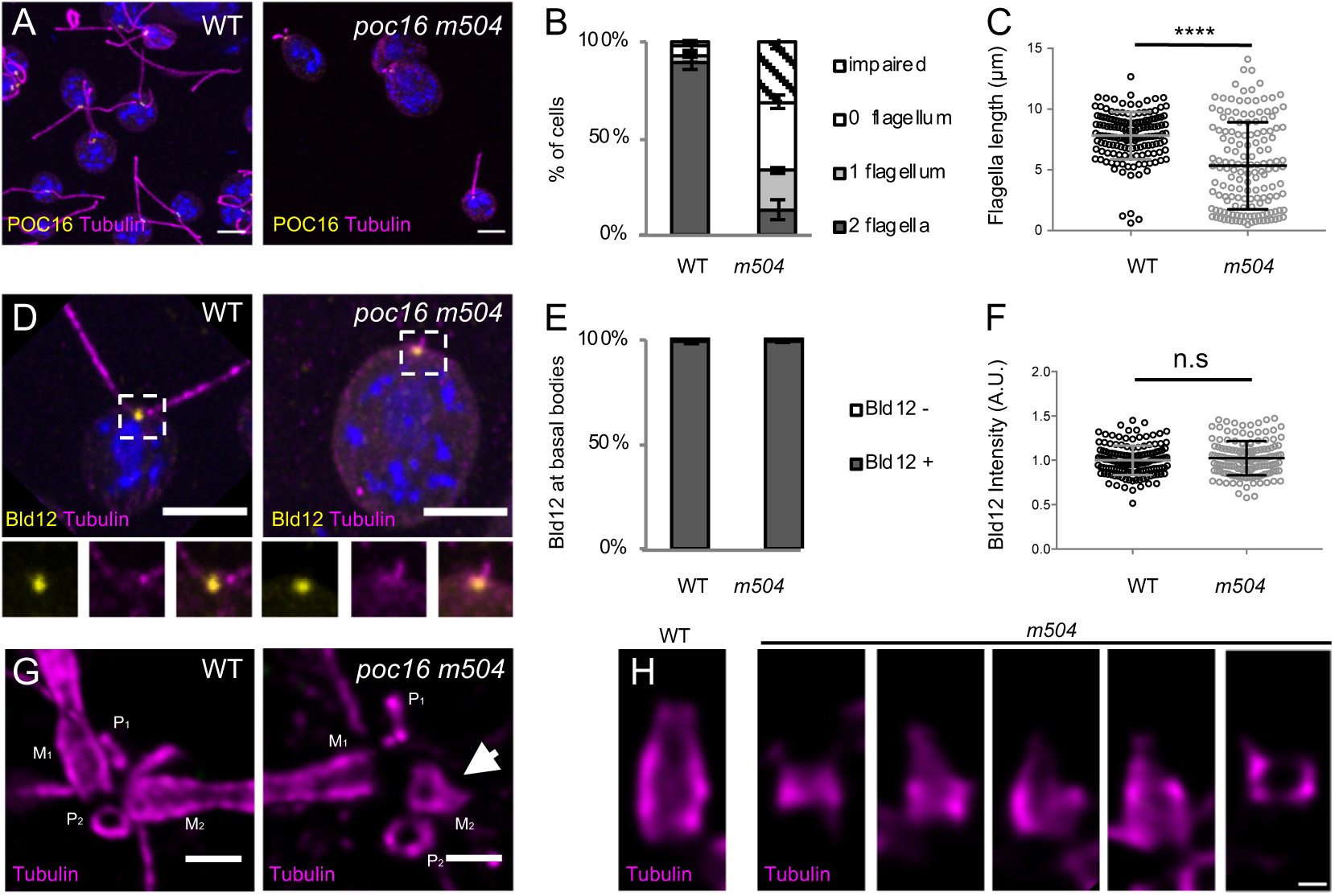
**POC16 is important for flagella assembly and proper centriolar structure.** (A) Confocal image of *Chlamydomonas* WT and *poc16 m504* cells stained with tubulin (magenta) and POC16 (yellow). Scale bar: 5µm (B) Percentage of cells with 0, 1, 2 or impaired flagella, n=300 cells/condition from 3 independent experiments. Average +/- SD: WT: 2 flagella= 95.5% +/- 4, 1 flagellum= 2.2% +/- 2, no flagellum= 2.3% +/- 2, impaired flagellum= 0% +/- 0; *poc16 m504:* 2 flagella= 13% +/- 5, 1 flagellum= 20% +/- 2, no flagellum: 13% +/- 3, impaired flagellum= 31% +/-3. (C) Flagellar length in µm, n=150 cells/condition from 3 independent experiments. Mann Whitney test p<0.0001. (D) *Chlamydomonas* WT and *poc16 m504*-mutant stained with tubulin (magenta) and Bld12 (yellow). Scale bar: 5µm (E) Percentage of cells positive for Bld12 at centrioles, n=300 cells/condition from 3 independent experiments. Average +/- SD: WT Bld12 positive (Bld12+) 99% +/- 1, *poc16m504*: 99% +/- 1. Normality assessed by Pearson test, Welch T test p< 0.0001. (F) Bld12 fluorescence intensity, n=150 cells/condition from 3 independent experiments. Average +/- SD: WT= 1 +/- 0.01 (A.U.), *poc16m504*= 1 +/- 0.02 (A.U.). Normality assessed by Pearson test, Welch T test p=0.0714. (G) *In cellulo Chlamydomonas* WT or *poc16m504* centrioles/flagella U-ExM expanded stained for tubulin (magenta). M stands for mature centriole, P for procentriole. Arrows point to defective mature centriole. Scale bar: 400nm (H) Gallery of *poc16m504* defective short mature centrioles stained with tubulin (magenta) compared to a WT mature centriole (left panel). Scale bar: 100nm.

**Figure S7.**
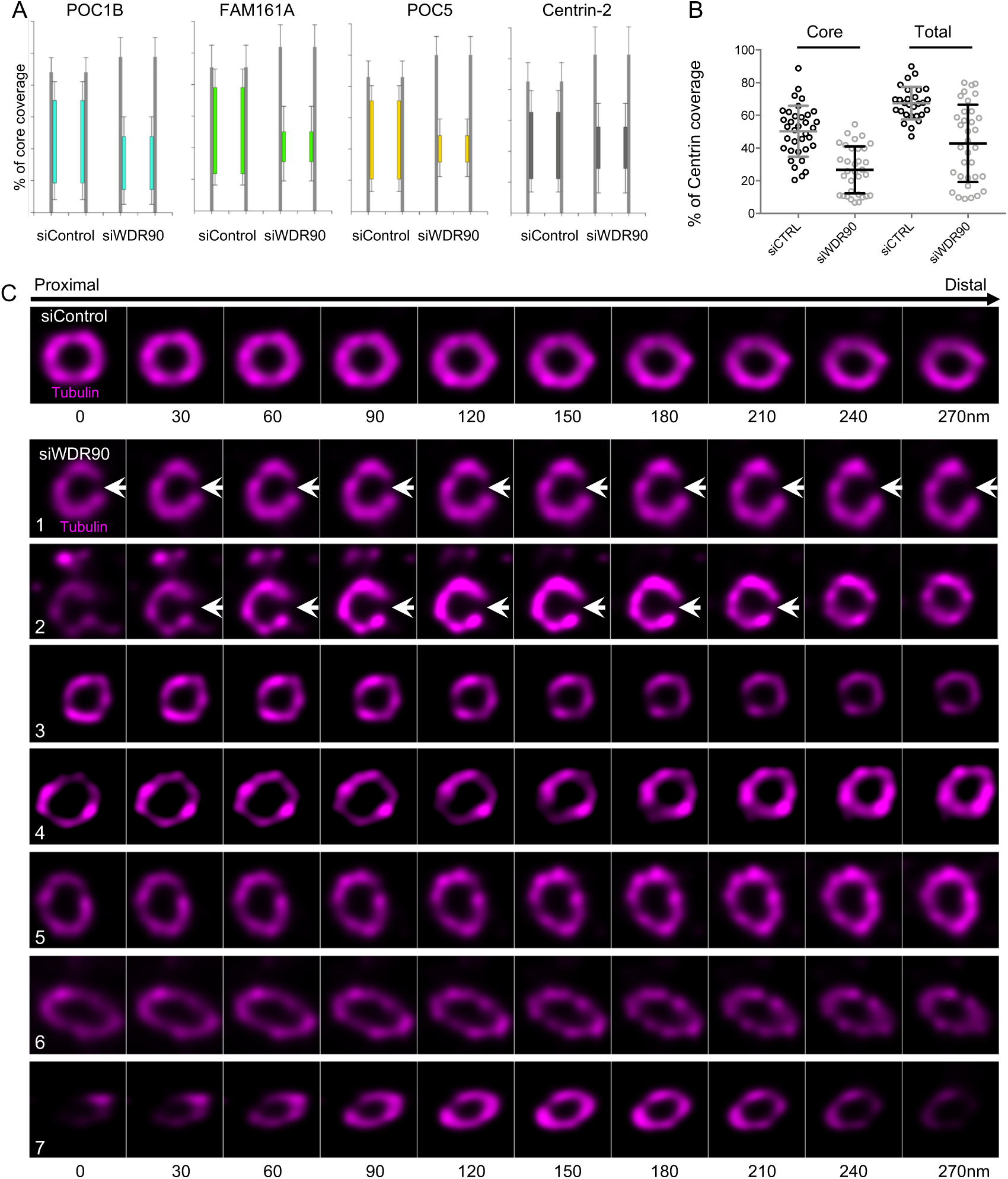
**WDR90 depletion leads to severe centriolar structure defects.** (A) Inner scaffold protein length based on Figure 5D and 5E, n>30 centrioles/condition from 3 independent experiments. (B) Centrin-2 length based on Figure 5D, measuring inner core or total (core + distal) length. (C) *In cellulo* U-ExM expanded centriole from U2OS cells treated with siRNA targeting scramble genes or *wdr90* stained for tubulin, top views. White arrows indicate centriole fracture. Scale bar: 200nm

**Video1. U-ExM expanded control centrioles.**

Top viewed *in cellulo* U-ExM expanded centriole from U2OS cell treated with scramble siRNA and stained for tubulin (magenta) and POC5 (green). Z-stack acquired every 0.12µm from the proximal to distal end of the centriole.

**Video2. U-ExM expanded centrioles depleted of WDR90.**

Top viewed *in cellulo* U-ExM expanded centriole from U2OS cell treated with *wdr90* siRNA and stained for tubulin (magenta) and POC5 (green). Z-stack acquired every 0.12µm from the proximal to distal end of the centriole.

## Supplementary Tables

**Table 1:** Tubulin and inner scaffold proteins fluorescence intensity on microtubule triplets from external to internal.

| x value for maximal intensity peak (nm) | Tubulin | Inner scaffold proteins |
| --- | --- | --- |
| **POC1B** | 53.99 | 68.48 |
| **FAM161A** | 51.36 | 73.75 |
| **POC5** | 46.09 | 72.43 |
| **Centrin-2** | 47.41 | 77.70 |

**Table 2:**
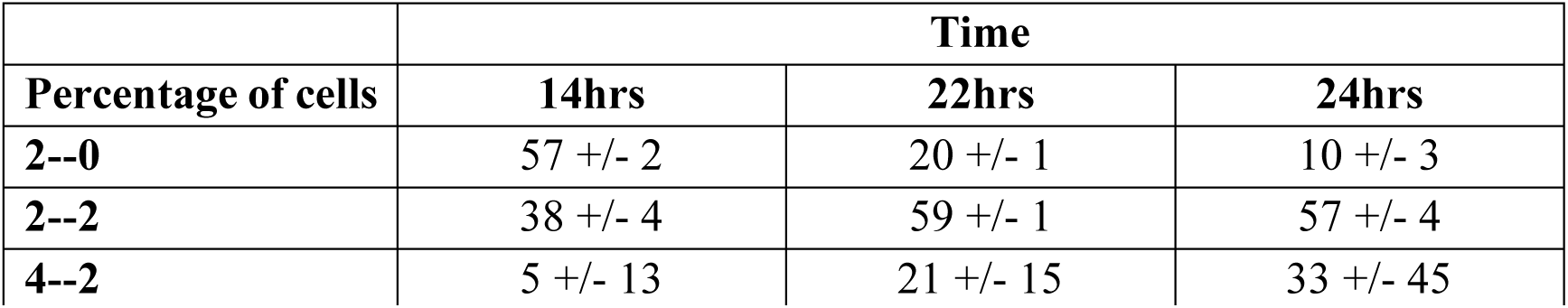
Percentage of cells with the following number of dots/cell respectively for WDR90 and Centrin-2.

|  | Time |  |  |
| --- | --- | --- | --- |
| Percentage of cells | 14hrs | 22hrs | 24hrs |
| **2--0** | 57 +/- 2 | 20 +/- 1 | 10 +/- 3 |
| **2--2** | 38 +/- 4 | 59 +/- 1 | 57 +/- 4 |
| **4--2** | 5 +/- 13 | 21 +/- 15 | 33 +/- 45 |

**Table 3:** Percentage of cells with the following number of dots/cell respectively for WDR90 and HsSAS-6.

|  | Time |  |  |
| --- | --- | --- | --- |
| Percentage of cells | 14hrs | 22hrs | 24hrs |
| **2--0** | 53 +/-3 | 19 +/- 6 | 9 +/- 3 |
| **2--2** | 44 +/- 4 | 67 +/- 1 | 67 +/- 3 |
| **4--2** | 3 +/- 1 | 13 +/- 1 | 24 +/- 4 |

**Table 4:** Diameter at proximal, core and distal region of the centriole

| Diameter | siControl | siWDR90 |
| --- | --- | --- |
| **Proximal** | 243 +/-23 nm | 231 +/- 16 nm |
| **Core** | 244 +/-25 nm | 223 +/- 15 nm |
| **Distal** | 214 +/- 25 nm | 203 +/- 15 nm |

**Table 5:** Inner scaffold proteins length coverage

| Length coverage | siControl | siWDR90 |
| --- | --- | --- |
| **POC1B** | 59 +/-14 nm | 34 +/- 16 nm |
| **FAM161A** | 61 +/- 9 nm | 18 +/- 7 nm |
| **POC5** | 56 +/- 11 nm | 17 +/- 9 nm |
| **Centrin-2** | 50 +/- 16 nm | 27 +/- 14 nm |

**Table 6:** Percentage of cells in each phase of the cell cycle according to post-mitotic time point

| Percentage of cells | Time |  |  |
| --- | --- | --- | --- |
|  | 14hrs | 22hrs | 24hrs |
| G0/G1 | 47 +/- 10 | 19 +/- 10 | 14 +/- 5 |
| S | 43 +/- 4 | 39 +/- 13 | 17 +/- 5 |
| G2/M | 10 +/- 1 | 42 +/- 7 | 69 +/- 5 |

**Table 7:** Percentage of cells displaying 0, 1, 2 or 4 dots of WDR90 based on the number of Centrin-2 dots in U2OS cells treated with siRNA targeting scramble genes or *wdr90*

| % of cells | 2 Centrin-2 dots |  |  | 4 Centrin-2 dots |  |  |
| --- | --- | --- | --- | --- | --- | --- |
|  | 0 WDR90 | 1 WDR90 | 2 WDR90 | 1 WDR90 | 2 WDR90 | 4 WDR90 |
| siControl | 0 +/- 0 | 10 +/- 2 | 90 +/- 2 | 0 +/- 0 | 71 +/- 1 | 29 +/- 1 |
| siWDR90 | 0 +/- 0 | 69 +/- 7 | 31 +/- 7 | 77 +/- 5 | 23 +/- 5 | 0 +/- 0 |

**Table 8:** Percentage of cells displaying 0, 1, 2 or 4 dots of POC5 based on the number of HsSas-6 dots in U2OS cells treated with siRNA targeting scramble genes or *wdr90*

| % of cells | 0 HsSAS-6 dot |  |  | 2 HsSas-6 dots |  |  |
| --- | --- | --- | --- | --- | --- | --- |
|  | 0 POC5 | 1 POC5 | 2 POC5 | 1 POC5 | 2 POC5 | 4 POC5 |
| siControl | 0 +/- 0 | 14 +/- 5 | 86 +/- 5 | 4 +/- 4 | 68 +/- 6 | 28 +/- 4 |
| siWDR90 | 0 +/- 0 | 61 +/- 10 | 39 +/- 10 | 63 +/- 9 | 29 +/- 11 | 8 +/- 3 |

## EXPERIMENTAL MODEL AND SUBJECT DETAILS

### *Chlamydomonas reinhardti* strains

*Chlamydomonas* strains control wild-type WT (cMJ030, *Chlamydomonas* Resource Center) as well as *poc16 m504* (LMJ.RY0402.069504, *Chlamydomonas* Resource Center) were described and cultured similarly to Hamel et al, 2017.

### Human cell lines

Human U2OS and RPE1 p53-cells (gift from Meng-Fu Bryan Tsou) were cultured similarly to Hamel et al, 2017. Cells were grown in DMEM supplemented with GlutaMAX (Life Technology), 10% tetracycline-negative fetal calf serum (life technology), penicillin and streptomycin (100 µg/ml).

To generate inducible episomal U2OS:GFP-WDR90RR cell line, U2OS cells were transfected using Lipofectamine 3000 (Life Technology). Transfected cells were selected for 6 days using 1µg/mL puromycin starting day 2 after transfection. Selected cells were amplified and frozen. For further experiments, U2OS:GFP-WDR90 cell line was grown in the medium specified above supplemented with 1µg/mL puromycin.

## METHOD DETAILS

### Ultrastructure Expansion Microscopy (U-ExM)

The following reagents were used in U-ExM experiments: formaldehyde (FA, 36.5-38%, F8775, SIGMA), acrylamide (AA, 40%, A4058, SIGMA), N,N’-methylenbisacrylamide (BIS, 2%, M1533, SIGMA), sodium acrylate (SA, 97-99%, 408220, SIGMA), ammonium persulfate (APS, 17874, ThermoFisher), tetramethylethylendiamine (TEMED, 17919, ThermoFisher), nuclease-free water (AM9937, Ambion-ThermoFisher) and poly-D-Lysine (A3890401, Gibco).

Monomer solution (MS) for one gel is composed of 25 μl of SA (stock solution at 38% (w/w) diluted with nuclease-free water), 12.5 μl of AA, 2.5 μl of BIS and 5 μl of 10X phosphate-buffered saline (PBS).

For isolated *Chlamydomonas* basal bodies (Klena et al., 2018), U-ExM was performed as previously described (Gambarotto et al., 2019). *In cellulo Chlamydomonas* U-ExM was performed on cells sedimented for 15 min on Poly-D-lysine-coated coverslips. Briefly, coverslips were incubated in 1% AA + 0.7% FA diluted in 1X PBS (1X AA/FA) for 5hrs at 37°C prior to gelation in MS supplemented with TEMED and APS (final concentration of 0.5%) for 1h at 37°C and denaturation for 30min at 95°C. Specifically, gels were stained for 3hrs at 37°C with primary antibodies against tubulin monobody (A345) (1:250 for cells and 1:500 for isolated basal bodies, scFv-F2C, Alpha-tubulin) (Nizak et al., 2003) and POC16 (1:100) (Hamel et al., 2017) or POB15 (1:100) (Hamel et al., 2017) diluted in 2% PBS/BSA. Gels were washed 3×10min in PBS with 0.1% Tween 20 (PBST) prior to secondary antibodies incubation for 3hrs at 37°C and 3×10min washes in PBST. A second round of expansion was done 3×150mL ddH20 before imaging.

Human U2OS cells were grown on 12mm coverslips and processed as previously described (Le Guennec et al., 2020). Briefly, coverslips were incubated for 5 hours in 2% AA + 1.4% FA diluted in 1X PBS (2X AA/FA) at 37°C. Denaturation was performed for 1h30 at 95°C and gels were stained as described above. The following primary antibodies were used: tubulin monobodies AA344 (1:250, scFv-S11B, Beta-tubulin) and AA345 (1:250, scFv-F2C, Alpha-tubulin) (Nizak et al., 2003), rabbit polyclonal anti-POC1B (1:250, PA5-24495, ThermoFisher), rabbit polyclonal anti-POC5 (1:250, A303-341A, Bethyl), rabbit polyclonal anti-FAM161A (1:250) (Le Guennec et al., 2020), mouse monoclonal anti-Centrin-2 (1:250, clone 20H5, 04-1624, Merck Millipore), rabbit polyclonal anti-WDR90 (1:250, NovusBio NBP2-31888). Specifically, staining against WDR90 was performed overnight at 37°C.

The following secondary antibodies were used: goat anti-rabbit Alexa Fluor 488 IgG H+L (1:400, A11008) and goat anti-mouse Alexa Fluor 568 IgG H+L (1:250, A11004) (Invitrogen, ThermoFisher).

For each gel, a caliper was used to accurately measure its expanded size (Ex_size_ in mm). The gel expansion factor (X factor) was obtained by dividing Ex_size_ by 12mm, which corresponds to the size of the coverslips use for sample seeding.

Thus, X factor = Ex_size_ (mm)/12(mm). The table below shows the Ex_size_ and X factor for all the gels used in this study.

| **Gel** | **siControl**<br>**$Ex_{size}$ (X factor)** | **siWDR90**<br>**$Ex_{size}$ (X factor)** |
| --- | --- | --- |
| **POC1B (n=1)** | 53 mm (4.42) | 52 mm (4.33) |
| **POC1B (n=2)** | 49 mm (4.08) | 50.5 mm (4.21) |
| **POC1B (n=3)** | 50.5 mm (4.21) | 50.5 mm (4.21) |
| **FAM161A (n=1)** | 50 mm (4.16) | 50 mm (4.16) |
| **FAM161A (n=2)** | 50 mm (4.16) | 51 mm (4.25) |
| **FAM161A (n=3)** | 50 mm (4.16) | 50 mm (4.16) |
| **POC5 (n=1)** | 51 mm (4.25) | 50.5 mm (4.21) |
| **POC5 (n=2)** | 50 mm (4.16) | 50 mm (4.16) |
| **POC5 (n=3)** | 50.5 mm (4.21) | 49 mm (4.08) |
| **Centrin-2 (n=1)** | 50 mm (4.16) | 50 mm (4.16) |
| **Centrin-2 (n=2)** | 50 mm (4.16) | 50 mm (4.16) |
| **Centrin-2 (n=3)** | 49 mm (4.08) | 49 mm (4.08) |

Pieces of gels were mounted on 24mm round precision coverslips (1.5H, 0117640, Marienfeld) coated with poly-D-lysine for imaging. Image acquisition was performed on an inverted Leica TCS SP8 microscope using a 63X 1.4NA oil objective with Lightening mode at max resolution, adaptive as ‘Strategy” and water as “Mounting medium” to generate deconvolved images. 3D stacks were acquired with 0.12µm z-intervals and an x, y pixel size of 35nm

### Cloning, and transient overexpression in Human cells

GFP-WDR90-N(1-225)RR and GFP-WDR90(FL)RR were cloned in the Gateway compatible vector pEBTet-eGFP-GW. Previously generated RNAi-resistant WDR90 DNA (Hamel et al, 2017) was used as template for PCR amplification. In brief, inserts were first subcloned in pENTR-Age-AGT using the restriction sites AgeI and XbaI. Second, a Gateway reaction was performed to generate the final expression plasmids pEBTet-GFP-WDR90-N(1-225)RR and pEBTer-GFP-WDR90(FL)RR, which were sequenced verified prior to transfection in human cells.

For transient expression, U2OS cells were transfected using Lipofectamine 3000 (Life Technology). Protein expression was induced using 1µg/mL doxycycline for 48 hours and cells were processed for immunofluorescence analysis.

Cloning of the GFP-WDR90 construct used in Figure 2 was done as follows: Human WDR90 was cloned by nested RT-PCR using total RNAs extracted from human RPE1 cells. Three different fragments corresponding to aa. 1-578, 579-1138, 1139-1748 of WDR90 (based on Genebank sequence NP_660337) were amplified and cloned separately using the pCR Blunt II Topo system (Thermo Fisher Scientific). The full coding sequence was then reconstituted in pCR Blunt II by two successive cloning steps using internal *Nru* I and *Sal* I, introduced in the PCR primers and designed in order not to modify WDR90 aa sequence. WDR90 coding sequence was then cloned into a modified pEGFP-C1 vector (Clontech) containing *Asc* I and *Pac* I restriction sites.

### Immunofluorescence in Human cells

Cells grown on a 15 mm glass coverslips (Menzel Glaser) were pre-extracted for 15sec in PBS supplemented with 0.5% triton prior to iced-cold methanol fixation for 7min. Cells were washed in PBS then incubated in 1% bovine serum albumin (BSA) in PBS-T with primary antibodies against WDR90 (1:250), Centrin-2 (1:500), HsSAS-6 (1:100), PCM1 (1:500), CP110 (1:500), GFP (1:500), mCherry (1:500) or tubulin (1:500). Coverslips were washed in PBS for 30min prior to incubation with secondary antibodies (1:1000) for 1 hour at room temperature, washed again for 30min in PBS and mounted in DAPCO mounting medium containing DAPI (Abcam).

Imaging was performed on a Zeiss LSM700 confocal microscope with a PlanApo 63x oil immersion objective (NA 1.4) and optical sections were acquired every 0.33 1m, then projected together using ImageJ.

### Cloning and protein purification

The constructs encompassing the predicted DUF667 domain of crPOC16 (Uniprot: A8JAN3), hsWDR90 (Uniprot: Q96KV7), drPOC16 (Uniprot: F1RA29), btPOC16 (Uniref: UPI000572B175), ptPOC16 (Uniprot: A0DK60), xtPOC16 (Uniref: UPI0008473371) and rnPOC16 (Uniref UPI0008473371) were cloned into a pET based expression vector via Gibson assembly (Gibson et al., 2009).

All recombinant proteins contained a N-terminal thioredoxin (TrxA) tag, used to enhance the expression level and the solubility of the target protein, followed by a 6xHis tag and a 3C cleavage site.

Protein expression was carried out in *E. coli* BL21 (DE3) competent cells grown in LB media at 37°C to OD_600_ = 0.6 and induced for 16h at 20°C with 1mM IPTG. Cells were subsequently resuspended in lysis buffer (50 mM Hepes pH 8, 500 mM NaCl, 10% v/v glycerol, 10 mM imidazole pH 8, 5 mM β-mercaptoethanol) supplemented with DNase I (Sigma), complete EDTA-free protease inhibitor cocktail (Roche) and lysed by sonication. The supernatant was clarified by centrifugation (18000 rpm, 4°C, 45 min), filtered and loaded onto a HisTrap HP 5 ml column (GE Healthcare). After extensive washes with wash buffer (50 mM Hepes pH 8, 500 mM NaCl, 10% v/v glycerol, 20 mM imidazole pH 8, 5 mM β-mercaptoethanol), the bound protein was eluted in the wash buffer supplemented with 400 mM imidazole. For crPOC16, hsWDR90, drPOC16 and xtPOC16, a 10 to 400 mM imidazole gradient was required to successfully detach the protein from the column.

The protein-containing fractions were pooled together and dialysed against the lysis buffer at 4°C for 48 hours in the presence of the 6xHis-3C protease. The tag-free protein was reapplied onto a HisTrap HP 5 ml column (GE Healthcare) to separate the cleaved product from the respective tags and potentially uncleaved protein. The processed proteins were concentrated and further purified by size exclusion chromatography (Superdex-75 16/60, GE Healthcare) in running buffer (20 mM Tris pH 7.5, 150 mM NaCl, 2 mM DTT). Protein were analysed by Coomasie stained SDS-PAGE and the protein-containing fractions were pooled, concentrated and flash-frozen for storage at −80°C. All protein concentrations were estimated by UV absorbance at 280 nm.

### Microtubule binding assay

Taxol-stabilized microtubules (MTs) were assembled in BRB80 buffer (80 mM PIPES-KOH pH6.8, 1 mM MgCl_2_, 1 mM EGTA) from pure bovine brain tubulin at 1 mg/mL (Centro de Investigaciones Biológicas, Madrid, Spain). 50 µL of stabilized MTs were incubated with 20µL of protein at 1 mg/mL for 2 hours at room temperature. After centrifugation on a taxol-glycerol cushion (8’000 rpm, 30°C, 20min) the supernatant and the pellet were analyzed by Coomasie stained SDS-PAGE gels. As a control, MTs alone and each protein alone were processed the same way.

### Tubulin binding assay

Tubulin at 10 µM was incubated with a slight molar ratio excess of each protein construct (around 15 µM) in MES buffer for 15 min on ice. After centrifugation at 13’000 x g at 4°C for 20 min, the supernatant and the pellet were analyzed by Coomasie stained SDS-PAGE.

### *In vitro* microtubules decoration and imaging

For simple decoration, Taxol-stabilized microtubules were nucleated as described (Schmidt-Cernohorska et al., 2019) and subsequently exposed to recombinant WDR90-N(1-225) in a 1:1 molar ratio for 30min at room temperature. Five µL of protein complexes solution were blotted on Lacey carbon grids and stained with Uranyl Acetate (2%) for 3 then 30 seconds.

For double decoration, *in vitro* microtubules were incubated with WDR90-N(1-225) in a 1:1 molar ratio for 5min at room temperature prior to addition of 2X free tubulin for 30min at room temperature. Negatively stained grids were prepared as above. Similarly, double decorated microtubules were prepared for cryo-fixation.

Electron micrographs were acquired on a Technai 20 electron microscope (FEI Company) and analyzed using ImageJ.

### Mitotic shake off

RPE1 p53-cells were seeded in T300 flasks the day before shake off. Flasks were shaken vigorously to detach mitotic cells collected in medium. Cells were pelleted by centrifugation for 5min at 1000 rpm and suspended in 10nM EdU containing medium prior to seeding in 6 well plates onto 15mm coverslips. Cells were fixed at different time points and processed in parallel for immunofluorescence or FACS analysis.

### WDR90 depletion using siRNA

U2OS cells were plated onto 15mm coverslips in a 6-well plate and 10nM silencer select pre-designed siRNA s47097 was transfected using Lipofectamine RNAimax (Thermo Fischer Scientific). Medium was changed 4hrs and 48hrs post-transfection and cells were analyzed 96hrs post-transfection.

In U2OS:GFP-WDR90(FL-RR) stable cell line, RNA-resistant protein expression was induced constantly for 96hrs using 1µg/mL doxycycline.

### Immunofluorescence on *Chlamydomonas reinhardti* cells

*Chlamydomonas* cells were sediment on Poly-D-Lysine coated-12mm coverslips (Menzel Glaser) for 30min prior to 7min fixation in −20°C methanol. Cells were washed in PBS then incubated in 1% bovine serum albumin (BSA) in PBS-T with primary antibodies against POC16 (1:500), Bld12 (1:100) and Tubulin DM1*α* (1:500) for 1h at room temperature. Coverslips were washed in PBS for 30min prior to incubation with secondary antibodies for 1 hour at room temperature, washed again for 30min in PBS and mounted in DAPCO mounting medium containing DAPI (Abcam). Only isolated cells were analyzed, the rest of the cells, which were grouped in palmeloids were not analyzed.

Imaging was performed on a Zeiss LSM700 confocal microscope with a PlanApo 63x oil immersion objective (NA 1.4) and optical sections were acquired every 0.33 nm, then projected together using ImageJ.

### Electron microscopy on *Chlamydomonas reinhardtii*

For sample preparation, cells were pelleted for 5min at 500g, fixed in 2.5% glutaraldehyde/TAP 1X for 1h at RT and washed 3x in TAP 1X. Fixed cells were further treated with 2% osmium tetraoxyde in buffer and immersed in a solution of uranyl acetate 0.25% over night (Tandler reference) to enhance contrast of membranes. The pellets were deshydrated in increasing concentrations of ethanol followed by pure propylene oxide and embedded in Epon resin. Thin sections for electron microscopy were stained with uranyl acetate and lead citrate, and observed in a Technai 20 electron microscope (FEI Company).

Micrographs analyses were performed on ImageJ and GraphPadPrism7. Symmetrization on top views was performed using CentrioleJ pluggin (https://gonczy-lab.epfl.ch/resources/ressources-centriolej/). The UnwarpJ plugin was required to perform image circularization using the center of the nine A-microtubules as landmark points. A 9-fold symmetrization was then applied to the circularized image. The deformation parameters were adjusted depending on the quality of the original image. For WT: initial deformation à fine, final deformation à fine. For m504: initial deformationà fine, final deformationà very fine.

### Image analysis

For centrioles counting, immunofluorescences were analyzed on a Leica epifluorescence microcoscope.

For fluorescence intensity, maximal projections were used.

Confocal centrosomal intensities were assessed using an area of 20 pixels on Fiji. For each experiment, control values were averaged and all individual measures for control and treated conditions were normalized accordingly to obtain the relative intensity (A.U.). Normalized individual values were plotted on GraphPadPrism7.

Confocal centriolar intensities were assessed by individual plot profil (25 points) on each pair of mature centrioles. For each experiment, the average (Av) of control values was calculated and all individual measures for control and treated conditions were normalized on Av to obtain the relative intensity (A.U.). An average of all normalized measures was generated and plotted in GraphPadPrism7.

For U-ExM data, length coverage quantification was performed as previously published in (Le Guennec et al., 2020).

For top views, a measurement from the exterior to the interior of the centriole was performed on each microtubule triplet displaying a resolved signal for both tubulin and the core protein. For each tubulin measurement, the position (x-value) of the maximal fluorescence intensity of the core protein was aligned individually to the position of the respective tubulin maximal intensity. All individual values of distance were plotted and analyzed in GraphPadPrism7.

Measurements of diameter in siControl and siWDR90 conditions were performed on S-phase mature centrioles imaged in lateral view. Briefly, lines of 50 pixels thickness were drawn within the proximal, central and distal regions defined in respect with the position of inner core proteins POC5 and FAM161A. Proximal region was then defined as the portion of the centriole below staining of POC5 or FAM161A and the distal region as above. In the siWDR90 condition, proximal region was defined as below the remaining belt of POC5 of FAM161A, the core region was measured just above the remaining belt and the distal region as the last 100 nm of the centriole. The Fiji plot profile tool was used to obtain the fluorescence intensity profile from proximal to distal for tubulin and the core protein from the same line scan.

Roundness was calculated on perfectly imaged top views of centrioles by connecting tubulin peaks on ImageJ.

### Statistical analysis

The comparison of two groups was performed using a two-sided Student’s t-test or its non parametric correspondent, the Mann-Whitney test, if normality was not granted either because not checked (n < 10) or because rejected (D’Agostino and Pearson test). The comparisons of more than two groups were made using one or two ways ANOVAs followed by post-hoc tests (Holm Sidak’s or Sidak’s) to identify all the significant group differences. N indicates independent biological replicates from distinct sample. Every experiment, except for resin electron microscopy, was performed at least 3 times independently. Data are all represented as scatter or aligned dot plot with centerline as mean, except for percentages quantifications, which are represented as histogram bars. The graphs with error bars indicate 1 SD (+/-) and the significance level is denoted as usual (*p<0.05, **p<0.01, ***p<0.001, ****p<0.0001). All the statistical analyses were performed using Excel or Prism7 (Graphpad version 7.0a, April 2, 2016).

## Supplemental methods

### Protein alignment

The protein sequences were aligned using Clustal Omega and the secondary structure elements were predicted using Phyre 2, PONDR and XtalPred-RF.

### 3D model

The *Chlamydomonas* POC16 model was prepared using *Phyre2* (Kelley 2015 Nature Protocols) and refined against the FAP20 cryo-EM map EMD_20858 using *phenix.real_space_refine* (Afonine 2018 ActaD). Superposition of the POC16 model excluding flexible loops against FAP20 was done using *COOT* (Emsley 2010 ActaD) and yielded a Root Mean Square Deviation (RMSD) value of 1.6 Angs. The figures were prepared using *ChimeraX* (Goddard 2018 Protein Science).

### PtPOC16 antibody purification

To generate anti-PtPOC16 antibody, a fragment encoding amino acids 2-210 was used for rabbit immunization (Eurogentec). Antibodies were subsequently affinity-purified over a column of PtPOC16(2-210) immobilized on Affi-Gel 10 (Bio-Rad Laboratories) and dialyzed against PBS/5% glycerol.

### Immunofluorescence in *Paramecium tetraurelia*

Immunofluorescence was performed according to (Beisson et al., 2010). Briefly, Paramecia were permeabilized for 5min in 0.5% saponin in PHEM Buffer (PIPES 60mM, HEPES 25mM, EGTA 10mM, 2mM MgCl2 pH 6.9) and fixed in 2% paraformaldehyde (PFA) for 10 min. Cells were washed 3×10min in PHEM-saponin buffer and stained with primary antibodies against POC16 (1:50) and tubulin 1D5 (1:10) for 30min at room temperature. Cells were incubated with secondary antibodies for 20min, washed twice in PHEM-saponin prior to a last wash in TBST-BSA supplemented with Hoechst 2mg/mL.

Imaging was performed on a Zeiss LSM700 confocal microscope with a PlanApo 40x oil immersion objective (NA 1.4) and optical sections were acquired every 0.33 1m, then projected together using ImageJ.

### Human cells cold shock treatment

U2OS cells grown on 15mm coverslips and transiently overexpressing mCherry-WDR90-N(1-225)RR for 24hrs were placed in 4°C PBS for an hour on ice and fixed in −20°C methanol. Coverslips were processed for immunofluorescence using primary antibodies against mCherry (1:500) and anti-tubulin DM1α (1:1000).

### *In vitro* crPOC16 microtubule decoration

*In vitro* stabilized Taxol-microtubules were prepared in MES-BRB80 derived buffer in contrast to K-PIPES-BRB80 to allow crPOC16(1-295) protein solubility. Samples were then processed similarly to WDR90-N(1-225).

### FACS analysis

Cells were processed similarly to Macheret et al 2018. Post-mitotic cells were washed 2x with PBS then permeabilized and treated with Click-EdU-Alexa 647 (Carl Roth EdU Click FC-647, ref 7783.1) according to manufacturer’s instruction. Genomic DNA was stained with propidium iodide (Sigma, Cat. No. 81845) in combination with RNase (Roche, Cat. No. 11119915001). EdU-DNA content profiles were acquired by flow cytometry (Gallios, Beckman Coulter) to assess the percentage of cells that entered S phase in each condition at each time point.

### PCM1 depletion using siRNA

Stable inducible GFP-WDR90 U2OS cells were plated in doxycycline containing– medium onto 15mm coverslips in a 6 well plate and 20nM silencer select pre-designed siRNA ADCSU9L was transfected using Lipofectamine RNAimax (Thermo Fischer Scientific). Medium was changed 4hrs post-transfection and cells were analyzed 48 hours post-transfection.

### Contact for reagent and resource sharing

Further information and requests for resources and reagents should be directed to and will be fulfilled by the Lead Contact, Virginie Hamel.

